# Precision functional imaging in infants using multi-echo fMRI at 7T

**DOI:** 10.1101/2025.11.09.687453

**Authors:** Julia Moser, Alireza Sadeghi-Tarakameh, Julian S. B. Ramirez, Thomas J. Madison, Lana Hantzsch, Kimberly B. Weldon, Han H. N. Pham, Jacob T. Lundquist, Sooyeon Sung, Inyene Ukpong, Emily Drexler, Alyssa K. Labonte, Edward J. Auerbach, Gregor Adriany, Chad M. Sylvester, Steven M. Nelson, Yigitcan Eryaman, Jed T. Elison, Damien A. Fair, Essa Yacoub

**Affiliations:** Masonic Institute for the Developing Brain, University of Minnesota, Minneapolis, MN, USA; Center for Magnetic Resonance Research (CMRR), University of Minnesota, Minneapolis, MN, USA; Graduate School of Education, Kyung Hee University, Seoul, Republic of Korea; Department of Psychiatry, Washington University in St. Louis, St. Louis, MO, USA; Department of Pediatrics, University of Minnesota, Minneapolis, MN, USA; Institute of Child Development, University of Minnesota, Minneapolis, MN, USA

**Author notes:** These authors have contributed equally to this work (Damien A. Fair, Essa Yacoub).

## Abstract

Personalized functional brain developmental trajectories can be studied with Precision Functional Mapping (PFM). Our previous work has demonstrated that PFM can be achieved in infants despite rapid brain growth. However, even with extensive data collection (up to 1 hour of fMRI), the reliability and precision of these maps remain lower than those observed in youth and adults - particularly within subcortical structures. In this work we demonstrate the utility of high-field 7T MRI compared to 3T MRI for facilitating PFM in infants. We showcase data from multi-echo fMRI acquisitions in the same infants at both 7T and 3T and demonstrate that 7T imaging in infants is safe and feasible with our subject-specific safety workflow. Moreover, we demonstrate that the use of a higher magnetic field strength affords a spatial resolution more appropriately matched to infants’ smaller head and brain sizes, yielding notable improvements in data quality, especially for PFM. The increase in both spatial precision and reliability also suggests that 7T MRI can reduce the amount of data required for PFM. Last, we show how ultra-high field imaging can help us study the development of subcortical-to-cortical connectivity patterns, crucial for understanding brain development during this developmental window. 7T MRI is a promising new avenue for developmental cognitive neuroscience.

## Introduction

The human brain develops rapidly in both structure and function in the first 1000 days of life. Thereby, infants show significant individual level variability in brain functional architecture^1–3^. Personalized trajectories can be uncovered with Precision Functional Mapping (PFM) with functional magnetic resonance imaging (fMRI), which is a method to investigate brain functional architecture on an individual level, which can be informative about healthy development^4^. PFM traditionally requires the acquisition of large amounts of high-quality data from single individuals, to be able to precisely and reliably map brain functional organization, including small individual specific networks parts. In adults, depending on the conditions (smoothing, parcelations, acquisition, processing; see^5–7^) at least 45 minutes up to several hours of data are required to obtain a reliable map within an individual. Despite this burden, this paradigm shift in data acquisition has yielded several major insights into variations in individual functional cortical maps^5,8–10^

PFM in infants is especially challenging as the collection of large amounts of low motion data (typically during natural sleep) requires specialized acquisition strategies to maximize an infant’s comfort in the scanner environment^11^. At the same time, in order to concatenate data, data need to be acquired in a short period of time (e.g. subsequent days), as infant brains show rapid developmental shifts in brain anatomy and physiology, resulting in both structural^12^ and functional^13,14^ changes. As a consequence, PFM studies in very young individuals have been rare^3,15^. Further complicating things, the overall signal-to-noise ratios (SNR) are lower and partial volume effects are increased in infants^16^ when employing readily available hardware and acquisition protocols optimized for adults, due to infants’ small brain sizes. As such prior work suggests that the times typically acquired for PFM using a 3T scanner in adults are insufficient for infants^2^.

Along with these challenges, the typical spatial resolution for data acquisition at 3T for adults is also unlikely to be optimal for infants. In the adult brain the average cortical thickness is around 2.6 mm^17–20^, while the fast changing infant brain has cortical thicknesses that range from about 1.6-2.0 mm (over the first couple of months of life)^17,21^. As such, the commonly used 2 mm voxel size for functional imaging likely leads to extensive partial voluming and is not ideal for extracting data from the cortical ribbon alone, as the large voxels include signals from areas outside the cortex. Subcortical structures increase their volume by 130% in the first year of life^22^, highlighting that the need for smaller voxel sizes does not only apply to the cortex.

Across all ages, subcortical structures exhibit the lowest SNR and are the most difficult to delineate using PFM. This is primarily due to their greater distance from the coil elements relative to cortical surface structures. Nonetheless, the development of subcortical regions and connectivity between subcortical regions and the cortex during the first 1,000 days of life is playing an important role in the development of brain functional networks^23,24^. These connections often go understudied due to the poor signals (e.g.^2,25^). In adults, subcortical partitions of cortical functional networks have been identified^26^ and their individual specificity demonstrated with PFM^27,28^. Similar work in early development is lacking thus far.

In the light of these challenges, ultra-high field 7T imaging can improve our ability to conduct PFM and revolutionize infant scanning by increasing SNR, improving resolutions, and reducing scan times^29–32^. To do this, several hurdles will need to be overcome, primarily ensuring safety of infant 7T acquisitions (see^33–36^). The most widely used 7T system, the Siemens Magnetom Terra, is FDA-approved only for subjects weighing more than 30 kg. Therefore, rigorous safety assessments must be conducted on-site in compliance with international guidelines, particularly regarding the radiofrequency (RF) energy deposition in the tissue by the transmit coil. It can be minimized by limiting multiple factors^37^, including head specific absorption rate (SAR) and peak 10g-averaged local SAR (pSAR_10g_). In addition to an initial report on 7T infant imaging^35^, Malik et al.^33^ recently conducted comprehensive numerical studies using neonate models at 7T, estimating head SAR and pSAR_10g_ limits for infants undergoing MRI with a commercially available head coil. Their results suggest infant imaging at ultra-high fields to be practically feasible.

This work demonstrates how we can leverage 7T MRI for precision functional imaging to study the developing infant brain. We present data from the same infants at both 7T and 3T, including our subject-specific safety assessments to ensure subject safety. The results highlight the advantages of using 7T for studying individual specific subcortical and cortical functional development. The functional connectivity examples derived from this sample containing within subject 3T and 7T data showcase how ultra-high field imaging might help us rethink the development of functional architecture, and the degree of ‘maturation’ we assign to infant networks.

## Results

### Safety of infant scanning on a clinical 7T MRI scanner

Multi-echo fMRI with infants in their first few weeks of life on a ultra-high field clinical 7T scanner is safe and feasible and results in T_2_^*^ weighted images with high tissue contrast (Suppl Fig 1). To inform our subject-specific workflow for safe data acquisition that adheres to SAR limits, we developed an EM simulation coil model to emulate a commercially-available 7T head coil (Nova Medical, Wilmington, MA). To verify the similarity between the RF fields generated by our simulated coil model and those of the commercial coil used in this study, we compared simulated and measured B_1_^+^ maps (Suppl. Figure 2). The results demonstrate good agreement, with a normalized root-mean-square error (NRMSE) of only 7%. Figure 1A illustrates the power budget in the transmit head coil in the presence of the infant model. At any input power level, 20% of the power was reflected due to under-loading caused by the small infant model. For the same reason, despite the inclusion of a local shield for the coil, 50% of the power was radiated. Consequently, only 30% of the input power was absorbed by the infant model.

**Figure 1.**
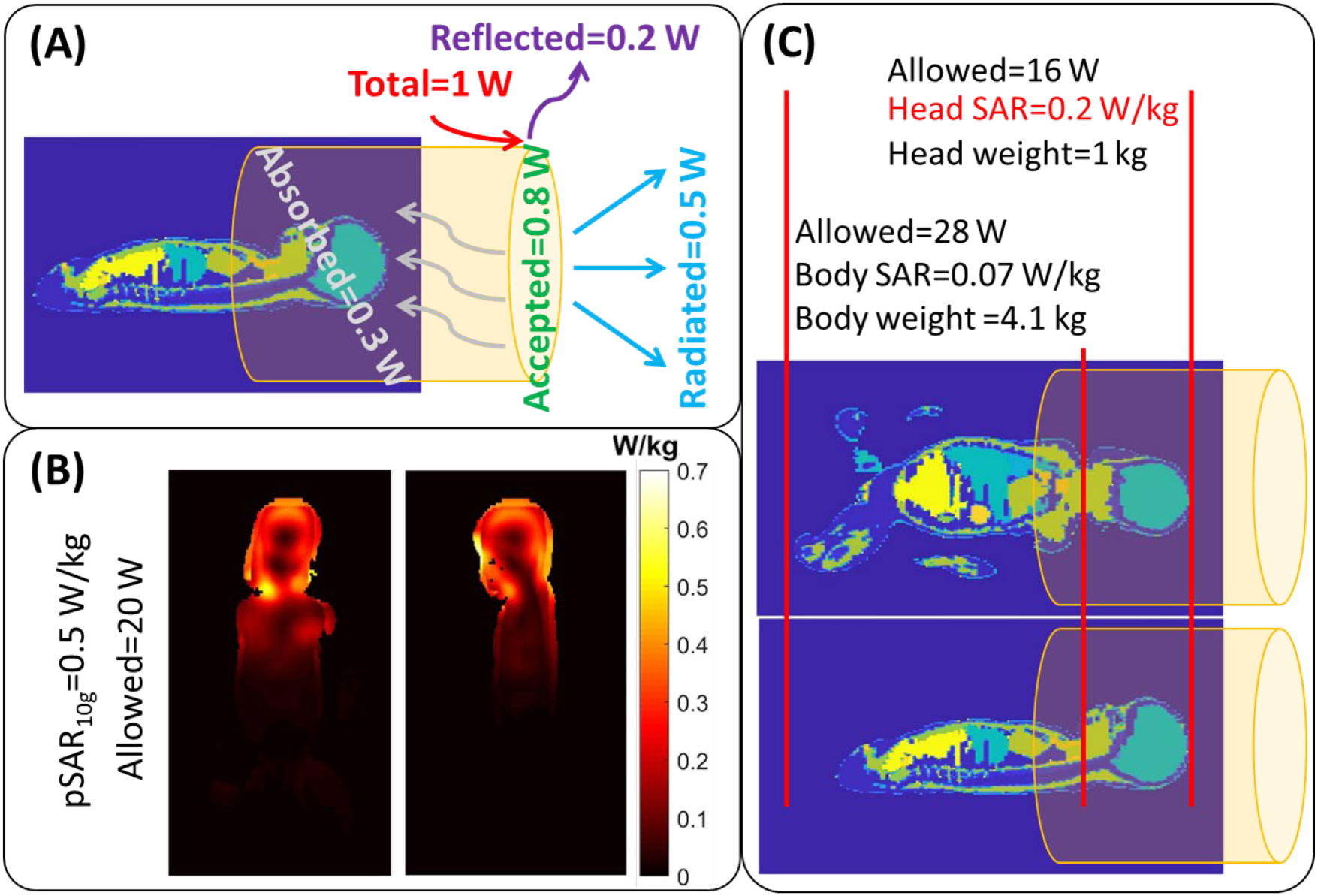
A) power budget in the transmit head coil in the presence of the infant model. B) 10g-averaged SAR map (peak SAR), along with C) head and body SAR calculations. Head SAR was determined as the limiting factor (head weight of 1 kg and head SAR of 0.2 W/kg (per 1 W input) results in a limit of 16 W to not exceed the head SAR guideline of 3.2 W/kg).

The peak 10 g-averaged local SAR (pSAR_10g_), head SAR, and body SAR calculations for this infant model are presented in Figures 1B & C. The maximum allowable power limits were calculated as 20 W, 16 W, and 28 W, corresponding to the thresholds defined by international safety guidelines ^37^ under normal operating mode for pSAR_10g_ (10 W/kg), head SAR (3.2 W/kg), and body SAR (2 W/kg), respectively. These results indicate that the head SAR threshold was reached before the local and body SAR thresholds, making head SAR the limiting factor in our infant safety pipeline. Based on this information, we developed an individual specific safety pipeline workflow to calculate the maximum reference voltage (*Vref*_*max*_), that can be used at the scanner to ensure scanning within safe head SAR levels (see Figure 9). *Vref*_*max*_ estimated using our safety pipeline workflow resulted in values similar to optimal (flip angle calibrated) values (*Vref*_*opt*_) calculated by the scanner, falling within the range of 175–200V. Our safety pipeline enabled straightforward verification of these values and adjustments when necessary.

We acquired on average 67 minutes of low motion data from each participant (6 infants, 5-12 weeks old) at 3T (PB015: 119 min; PB016: 75 min; PB017: 79 min; PB020: 76 min; PB021: 29 min; PB022: 26 min) and on average 30 minutes of low motion data at 7T (PB015: 17 min; PB016: 24 min; PB017: 16 min (10 min with 1.6 mm sequence and 6 min with 1.25 mm sequence); PB020: 41 min (26 min with 1.6 mm sequence and 15 min with 1.25 mm sequence); PB021: 27 min; PB022: 55 min (see Methods for details on data acquisition). The ratio of low motion data to total data acquired was comparable between 3T and 7T, indicating similar conditions for infants to comfortably sleep during the scan (PB015 89% at 3T and 82% at 7T; PB016 95% at 3T and 97% at 7T; PB017 91% at 3T and 80% at 7T; PB020 87% at 3T and 91% at 7T; PB021 73% at 3T and 99% at 7T; PB022 98% at 3T and 91% at 7T (eliminating two very high motion runs in each modality)).

The T_2_^*^ values calculated from the multi-echo acquisition replicated previous findings^38^ by showing shorter T_2_^*^ relaxation times at 7T compared to 3T (Figure 2, Suppl. Figure 3 and 4). The longer T_2_^*^ relaxation times previously found in 3T ME infant data^7^ was replicated in this sample (range of mean values 94-129 ms). The broad range of T_2_^*^ relaxation times previously reported in 3T infant data was observed across both field strengths. Figure 2A shows the distribution of T_2_^*^ relaxation times from cortical vertices in PB016 as an example.

**Figure 2.**
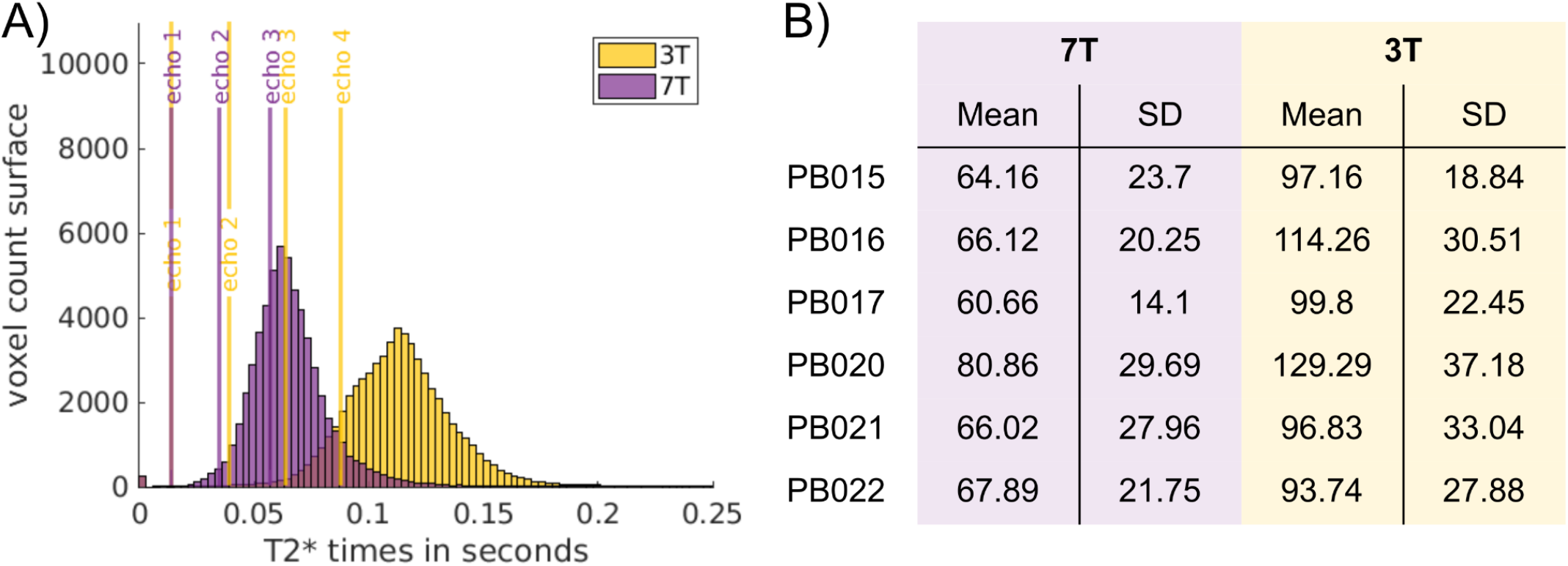
A) T_2_^*^ relaxation time distribution of cortical vertices for example participant PB016. Distributions for 3T and 7T are shown in relation to the acquired echoes. B) Mean and SD of T_2_^*^ relaxation times from cortical vertices for all participants.

### 7T fMRI data shows properties that are beneficial for PFM

#### An increase in spatial precision can be achieved with 7T

Higher resolution data acquired at 7T shows higher spatial precision as expected (3T data was acquired with 2 mm resolution, 7T data with 1.6 mm resolution). The full width at half maximum (FWHM) of the data in native space was ∼20% higher in 3T compared to 7T. The mean FWHM across runs across participants at 3T was 2.85 (SD = 0.24) and 2.36 (SD = 0.1) at 7T (Figure 3). This improvement in spatial precision was preserved throughout data processing and even persisted when data were normalized into adult MNI space (both 3T and 7T in 2 mm space). The mean FWHM across runs across participants in MNI space at 3T was 5.22 (SD = 0.32) and 4.32 (SD = 0.16) at 7T.

**Figure 3.**
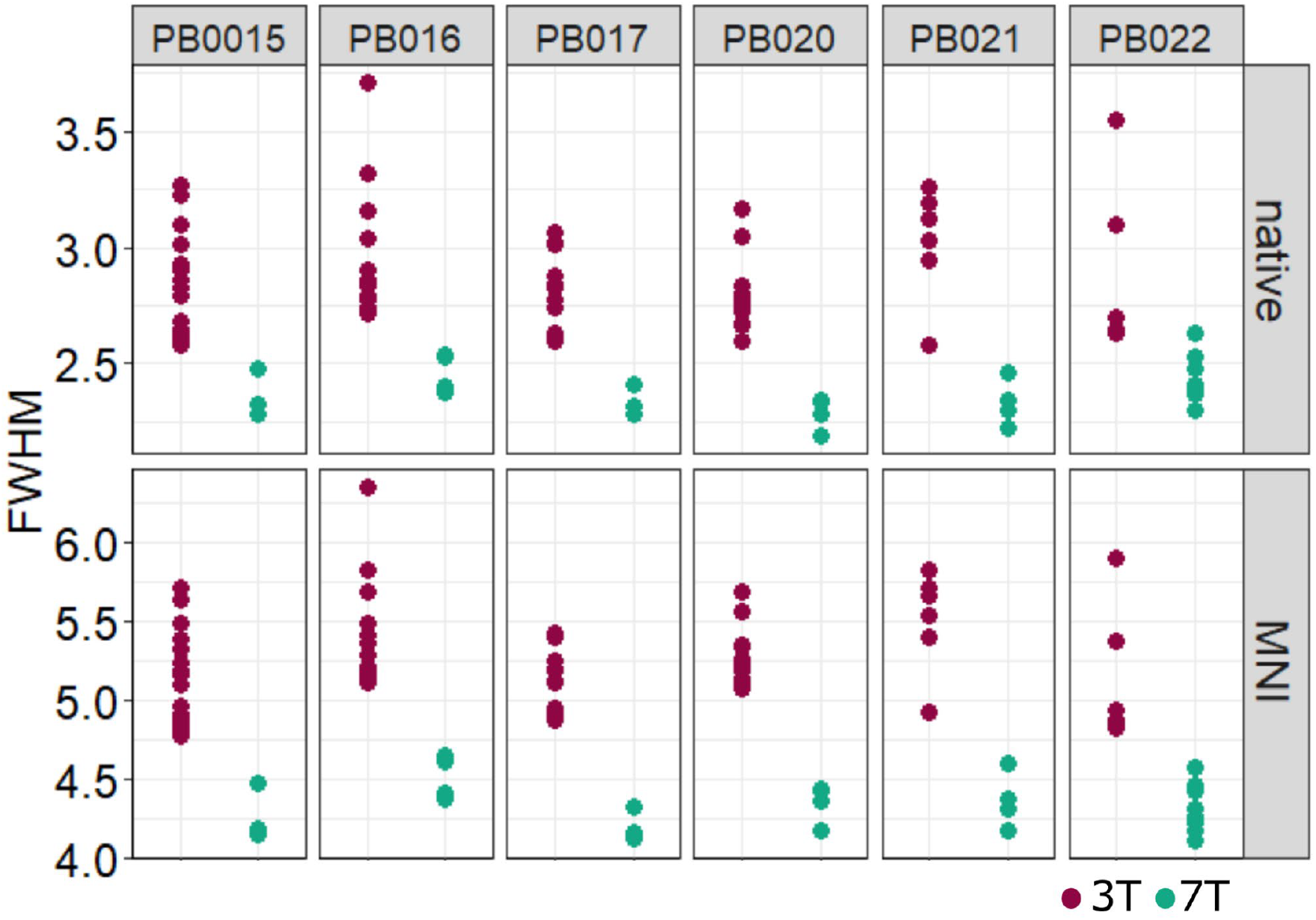
Full Width Half Max (FWHM) as measurement of spatial precision. Top: native space, Bottom: adult MNI space. Native space voxel size is 2 mm for 3T and 1.6 mm for 7T. MNI space voxel size is 2 mm for both. Each datapoint is one BOLD run of a given participant. Spatial precision is in both cases higher for 7T.

#### Functional connectivity calculated from 7T data shows more stability in less time

The stability of functional connectivity estimates is a key component of PFM. PFM is only possible with a reliable estimate of an individual’s functional architecture, which is traditionally achieved through the acquisition of large amounts of data. Data acquisition and processing choices can increase the stability of these estimates with shorter data amounts. We compared the stability of various amounts of data between 3T and 7T in PB016, PB020, PB021 and PB022; which were the subjects with more than 20 min of low motion data in both 3T and 7T acquisitions. Across the whole brain, maximal stability values (at 10 min) increased by 17-146% (PB016: 23%, PB020: 17%, PB021: 146%, PB022: 44%). Stability for cortical vertices increased by 15-120% (PB016: 15%, PB020: 21%, PB021: 120%, PB022: 18%) while for subcortical voxels, stability of functional connectivity estimates increased by 34-146% (PB016: 56%, PB020: 34%, PB021: 146%, PB022: 85%; Figure 4; Suppl. Figs. 5, 6, 7). The stability that could be achieved with 10 minutes of (2 mm) 3T data was reached with only 3-8 min of (1.6 mm) 7T data (PB016: 8, PB020: 8, PB021: 3, PB022: 6). This was true for correlations of cortical vertices (PB016: 8, PB020: 8, PB021: 3, PB022: 8), as well as subcortical voxels, for which this effect was even more pronounced (6 min of 7T data were approximately as stable as 10 min of 3T data for PB016, 7 min for PB020, 3 min for PB021 and 4 min for PB022).

**Figure 4.**
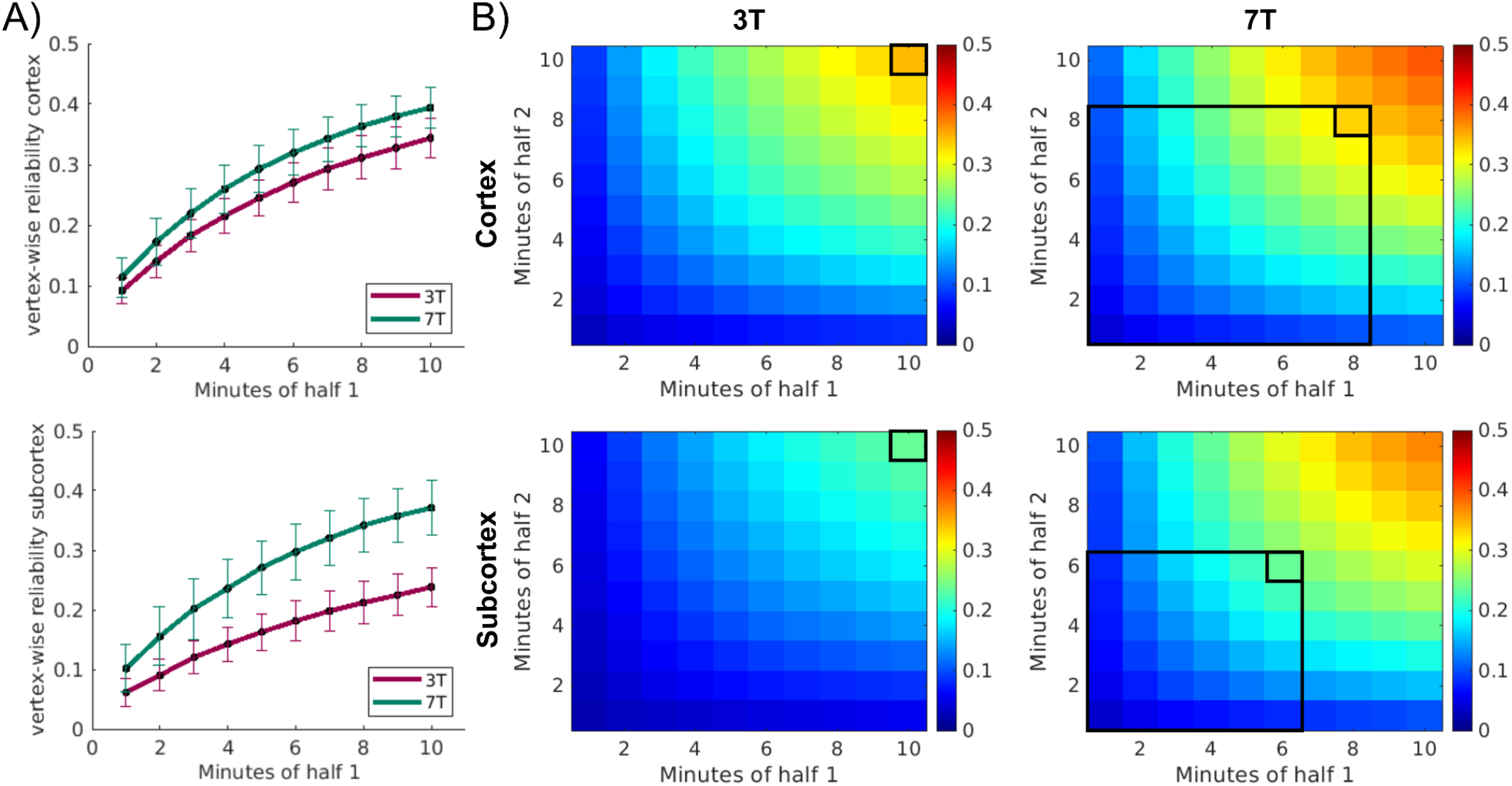
Stability heatmap PB016. A) reliability curves for cortical and subcortical vertices. Lines represent the average across vertices and permutations (100 permutations shuffling individual data minutes). Error bars represent the standard deviation across permutations. (Results correspond to the top row of heatmaps in B)) B) Each square represents the correlation of connectivity matrices when a set amount of data is used to create them (extending Figure A to also varying data amounts in half 2). Correlation strength is represented by the heatmap. Values are averaged across vertices and permutations. Rectangles indicate the amount of 7T data that is equivalent to the maximum in the 3T data.

This stability of functional connections in both 3T data and 7T data is not uniform across the brain. Figure 5 shows the spatial map of the upper right corner of the heatmap plot for PB016 (10 min vs. 10 min, see Suppl. Figures 8,9,10 for PB020-22). Notably, areas that already show to be relatively stable in 3T increase in 7T while areas with very low stability remain consistently low across acquisitions (e.g. inferior temporal and medial occipital cortex). All participants show this spatial trend, despite wide differences in the increase in stability in 7T. While the areas with the lowest stability are those tending to have the highest signal dropout, sensorimotor regions showed a relatively high stability.

**Figure 5.**
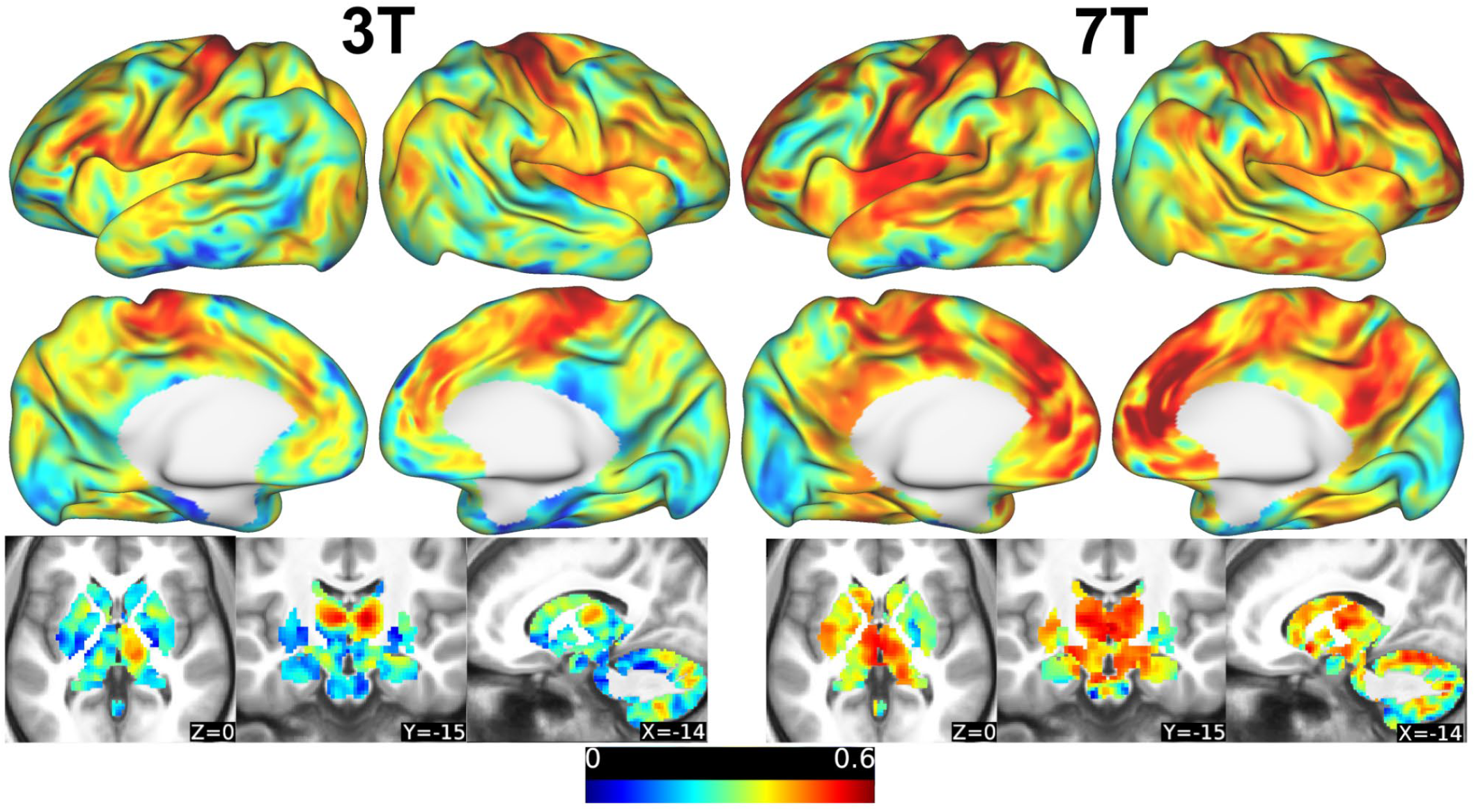
Spatial distribution of stability map from PB016 using 10 min vs. 10 min of low motion data (top right corner Figure 4 A and B) for 3T and 7T. Maps show common regions of low stability and regions of particularly high stability in 7T data.

#### Functional connectivity calculated from 7T data has stronger magnitudes

Low correlation magnitudes in functional connectivity matrices in infants complicate the detection of individual specific functional networks. Here we compare the correlation strength of functional connectivity matrices of individual participants between their 3T and 7T scans. All participants showed overall stronger or equivalent absolute correlation values in the 7T data compared to their 3T data (comparison of means: t(5) = 4.57, p =.006; Figure 6). This difference is particularly pronounced for connections that include subcortical vertices - within the subcortex and between the subcortex and other brain areas (Suppl. Figure 11). To ensure that this not only affects short range connectivity, we compared both correlation magnitudes for local connectivity (<1 cm) as well as for long range connectivity (>10 cm). Correlations are equivalent or stronger for both cases (Suppl Figure 12 & 13).

**Figure 6.**
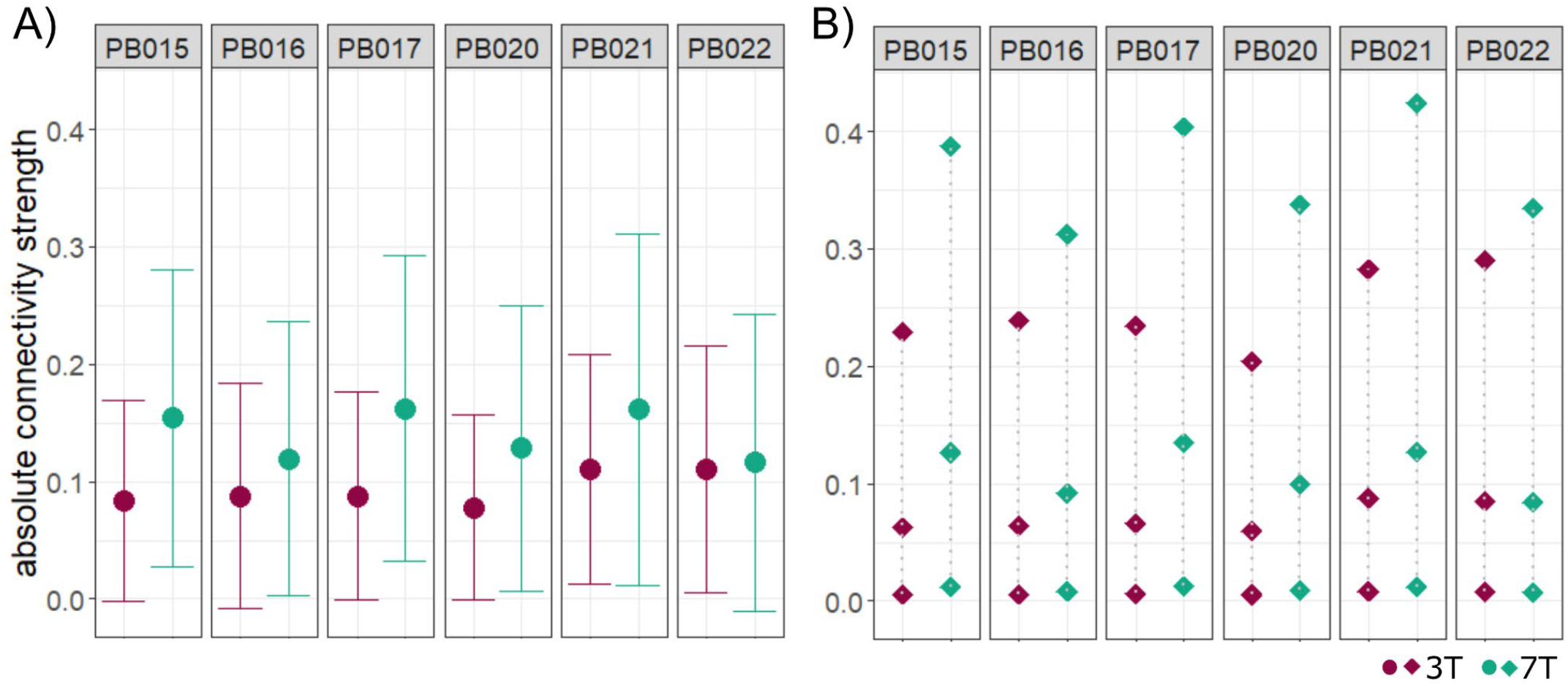
Absolute strength of functional connections compared between 3T and 7T data of each participant. A) mean with standard deviation of all absolute connections B) 5th, 50th and 95th percentiles of all absolute connections. All metrics show stronger or equivalent connectivity for 7T.

### Increase in resolution and connectivity strength shows real world impact

The differences between 3T and 7T acquisitions outlined above likely impact the investigation of functional networks with PFM in infants. Stronger and more spatially defined connections lead to more adult-like patterns of strong positive within network connectivity and negative connectivity with a variety of other networks (see Default Mode Network (DMN) as an example in Suppl. Figure 14). One of the most influential discoveries of PFM has been the identification of the Somato-Cognitive Action Network (SCAN)^39^, a network that interleaves effector specific regions in the motor cortex and which has upended the 90 year old model of motor cortex organization defined by the traditional homunculus^40^. This network shows individual differences and was believed to be missing in infants^39^. Figure 7 demonstrates the identification of the SCAN network within our PFM infant sample.

**Figure 7.**
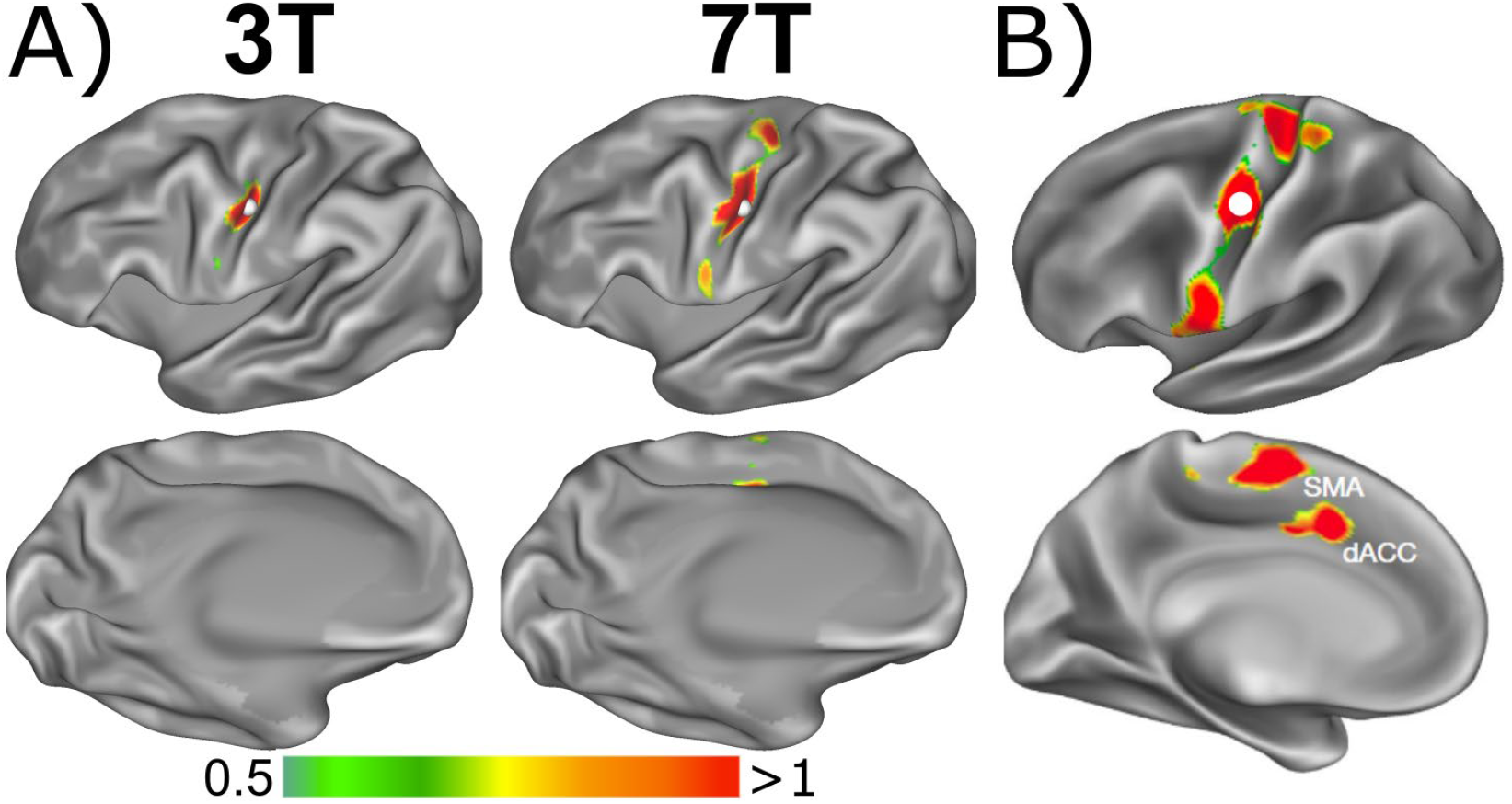
Use case examples for 7T improvements in PFM. A) Somato-Cognitive Action Network (SCAN) seed in PB020 (5 weeks old) using PB020’s 3T data and the 1.25 mm resolution 7T data. Increase in spatial resolution helps to delineate connectivity between SCAN hubs B) architecture of the adult SCAN - Figure adapted from Gordon et al. (2023)^39^.

We used seed-based connectivity within the motor strip to identify SCAN hubs. While the distinct three hub pattern of the SCAN is not very prominent in the 3T data, the increase in resolution with 7T helps to clearly differentiate the hubs, marking the distinct SCAN pattern. For this example, we leveraged the data acquired from PB020 with a 1.25 mm resolution, which is the highest spatial resolution we acquired in our sample. Findings replicated across other infants with 1.6 mm data, yet not as pronounced as with the 1.25 mm data (Suppl Figure 15).

## Discussion

The present study demonstrates that 7T imaging in infants is safe, feasible, and has distinct advantages for precision functional data acquisition compared to 3T imaging. The higher magnetic field strength allows for a spatial resolution more appropriate for infant’s small head and brain sizes, which reduces partial voluming effects and provides important benefits for data quality, particularly in the context of PFM.

### Practical considerations for safely imaging infants at 7T

For our safety assessment, we used a maximally conservative approach, being the first United States-based institution performing an ultra-high field fMRI study in infants. According to Figure 1A, only 30% of the total input power was dissipated in the infant model, indicating that a significant portion of the transmit power was either reflected or radiated rather than absorbed by the tissue. This highlights the under-loading effect caused by the small size of the infant head relative to the coil’s design. However, to introduce an additional safety buffer in the proposed workflow, a conservative approach was taken by using the total input power - rather than just the absorbed fraction - to calculate the head SAR. This ensures that even in the worst-case scenario, where power absorption may vary due to differences in individual anatomy or coil positioning, the SAR remains within safe limits. By adopting this precautionary method, the safety pipeline provides an extra margin of protection, mitigating potential risks associated with power distribution uncertainties in infant imaging at 7T. Despite using this very conservative workflow, we were able to run all the sequences within calculated power limits without compromising optimal flip angles. Further optimizations to the safety pipeline will increase automation to ensure that it is easily shareable with other research institutions using a Siemens 7T Terra scanner.

### 7T imaging increases feasibility of precision imaging in early development

The higher spatial resolution and the higher functional sensitivity that can be obtained with ultra-high field imaging greatly benefits precision functional imaging in infants. Our results show decreased spatial smoothness combined with stronger functional connections which help to more precisely isolate small individual network patterns. These improvements were apparent when the data were processed with standard processing pipelines optimized for 3T data. While using smaller voxels is entirely appropriate for the smaller infant brain, using smaller voxels at 3T would be more challenging and in the least would result in much longer scan times for PFM applications, based on the limited SNR that can be achieved in this scenario, which is not practical for infant studies.

In addition to spatial precision, reliability is a key element of precision functional imaging. Given that large amounts of data are needed to obtain reliable individual specific results in infants^3,7^, the increase in stability of functional connectivity we see with 7T will be critical to achieving reliability in more participants, as any reduction in scan time would be invaluable. The amounts of data gathered in these first participants did not allow for constructing conclusive reliability curves as have been done in previous studies^5,6^, with which it is possible to estimate a ceiling in reliability. But even a reduction from 10 to 8 minutes which was the minimal improvement in our stability analysis means an overall reduction of scan time by 20%. With the improvements afforded by 7T imaging, many different studies, such as those of patient populations who might not be able to participate in multiple consecutive visits, could benefit significantly from such higher quality precision style data.

Combining 7T infant MRI with other methodological advances, such as multi-echo (ME) fMRI^41^ and NORDIC thermal noise removal^42–44^ can further facilitate precision imaging in infants by increasing data reliability even more. The utility of these methods for 3T imaging in developmental populations has already been shown^7^. ME fMRI acquisitions, however, are less common at ultra-high fields due to the shorter T_2_^*^s compared to lower field strengths^45^. Importantly, though, the T2*s in infants are approximately 2-fold longer compared to adults^7,46,47^ making the combination of ME fMRI and ultra-high-fields a unique opportunity for infant applications.

### The SCAN network in neonates is discovered at 7T

As noted above, the SCAN network is a new discovery showing an interdigitated system within the motor cortex important for action control^39^. This discovery broke the 90 year understanding established by Penfield^40^ of how the motor cortex is organized. Gordon et al.^39^ showed the presence of the SCAN in older adults, children, and even older infants but not in data from newborns. Further work needs to investigate whether the SCAN is established shortly after birth to assess whether their findings can be attributed to the challenges in signal quality in 3T infant data. Leveraging the improvements with 7T demonstrated here, we clearly showed the presence of the SCAN network at a very young age (here 5 weeks). These data highlight the extremely early presence of this association network but also show how important our new methods are to ground interpretation of data collected at 3T.

Our results furthermore show that characteristic negative correlations (e.g. DMN with motor cortex) are strengthened in 7T data, contributing to the increase in absolute connectivity strength and giving connectivity patterns a more adult-like appearance. The increase in absolute connectivity strength in 7T data is furthermore not limited to short range connectivity but also applies to long-range connectivity (Suppl Figures 12 & 13) and specifically to connections with subcortical structures. Given previous findings of weaker anterior to posterior connections in infants^25^, 7T acquisitions could improve our ability to track the development of networks like the DMN, with important anterior to posterior hubs. Improving our ability to precisely characterize the development of functional brain architecture will be impactful for designing personalized interventions.

### Limitations and outlook

Given the benefits showcased here for performing functional imaging in populations with small brain structures, we predict 7T imaging to be an important tool for developmental cognitive neuroscience in the future. However, despite their clinical readiness, 7T scanners are not widely available, which will prevent the majority of scientists who do infant imaging to switch from 3T to 7T. Utilizing the increased precision from data acquired at 7T to develop models that can be used as priors in data analyses from 3T acquisitions will therefore also be a promising avenue for the field to benefit from infant 7T data. Refinement of data preprocessing methods specifically for infant 7T data will improve outputs further.

The variability of results in our small sample demonstrates that data acquisition at a higher field strength does not solve all the problems of infant imaging. Our stability analysis, for example, showed that 10 minutes of data are still not highly reliable in most cases when investigating functional connectivity. Similarly, motion is still a major confounding factor and further investigations need to be done regarding the impact of motion and how it might depend on field strength. As an example, PB0021, who showed the biggest improvements in stability between 3T and 7T also had the lowest percent of low motion data in their 3T data (73%) and the highest in their 7T data (99%), likely boosting this effect and making this subject less representative for the group.

In the present use case, we explored a three echo ME fMRI sequence to compare results between 3T and 7T. While ME fMRI is becoming more common in 3T acquisitions, it is still rare for 7T and non-existent at higher spatial resolutions. ME fMRI acquisitions in infants at 7T would benefit from further explorations to determine the ideal echo times for a given spatial resolution and to find the balance between accommodating the longer and broader T_2_^*^ relaxation times, while maximizing signal to noise for the higher spatial resolution. Future studies with larger samples in a broader age range will also be useful for determining optimal protocols.

This line of research is still in its infancy, with the results presented herein, a number of uncharted possibilities are already foreseeable for future studies. Safety procedures will need to be streamlined and generalized across different coils, including more optimal infant specific coils, and other sequences, such as anatomical or diffusion weighted imaging, will need to complement the fMRI data.

## Conclusions

7T imaging in infants is safe and feasible and our initial analyses show clear benefits for PFM over 3T imaging in an infant sample. A publicly available safety workflow will facilitate infant imaging at 7T at other institutions, enabling more precise studies on infant functional brain development. Improving our ability to study the development of subcortical-to-cortical and small cortical networks makes 7T MRI a promising tool for developmental neuroscience.

## Methods

### Sample

The sample for this study consists of six infants enrolled at the University of Minnesota. PB015 (48 weeks postmenstrual age (wPMA)), PB016 (46 wPMA), PB017 (51 wPMA), PB020 (45 wPMA), PB021 (46 wPMA) and PB022 (52 wPMA). Data included in this manuscript were collected in the framework of a study approved by the University of Minnesota’s Institutional Review Board, and written informed consent was obtained from both parents of the participants. Data for each participant were acquired over three to four days within one week (PB015: 3 days 3T, 1 day 7T; PB016: 2 days 3T, 1 day 7T; PB017: 2 days 3T, 1 day 7T; PB020: 2 days 3T, 1 day 7T; PB021: 2 days 3T, 1 day 7T; PB022: 2 days 3T, 2 days 7T). All data were acquired during natural sleep following established protocols^48^.

### Safety Assessment and Monitoring

#### Coil Model and Validation

Based on informed conjecture and existing literature^33,49^, an EM simulation coil model was developed to emulate a commercially-available 7T head transmit coil (Nova Medical, Wilmington, MA). Using CST Studio Suite (Darmstadt, Germany), EM simulations were conducted within a spherical phantom (Figure 8A) with ε_r_ = 50 and σ = 0.56 S/m. Both coil ports were matched to 50Ω better than −16 dB at 297 MHz. The simulation model was validated through experimental B_1_^+^ mapping within a similar spherical phantom using the Nova coil. The numerical and experimental B_1_^+^ maps were quantitatively compared using the NRMSE.

**Figure 8.**
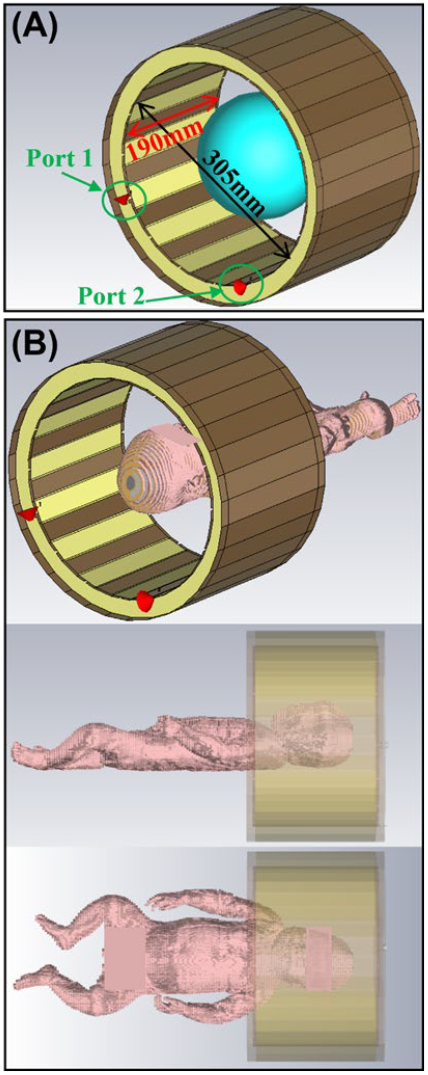
EM simulations with a spherical phantom (A) and an eight-week-old infant model (B).

#### Safe SAR Limits

The phantom was replaced with an eight-week-old infant voxel model (Figure 8B), available in CST Studio with a 1 kg head weight. To determine safe operating power limits, various parameters were monitored, including input, reflected, accepted, radiated, and tissue-absorbed powers, as well as peak 10g-averaged, head, and body SARs. Ensuring compliance with safety SAR limits^37^ - set at 10 W/kg for peak 10g-averaged SAR, 3.2 W/kg for head SAR, and 2 W/kg for body SAR - the study concluded that the head SAR limit was reached before the peak 10g-averaged SAR limit.

#### Subject-Specific Power Calculations

Based on this finding, a workflow (Figure 9) was proposed to ensure the safe operation of a 7T Siemens Magnetom Terra scanner for infant imaging:

1. First, the infant’s head weight (w) was estimated using a 3T anatomical scan (3D T1-weighted data). The maximum allowable input power (*P*_*max*_) of the coil was then calculated by multiplying this weight by the 3.2 W/kg head SAR limit suggested in international guidelines ^37^.
2. Next, an offline study was conducted using the full imaging protocol-including all pulse sequences intended for infant imaging - to identify the worst-case pulse sequence (WCPS), defined as the sequence with the highest power requirement. This was determined along with its power level 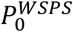 for a given reference voltage (*Vref*_*0*_) - both of which are typically reported in the log file of the scanner. In Siemens’s convention, the reference voltage denotes the transmitter voltage required to achieve a 180 ° flip angle using a 1 ms square pulse. Based on this analysis, a protocol with a fixed maximum reference voltage (*Vref*_*max*_) for each specific head weight was selected.
3. Finally, at the start of each in vivo session, Siemens’ routine power calibration step was performed to determine the optimal reference voltage (*Vref*_*opt*_). If the system’s power calibration suggested a *Vref*_*opt*_ exceeding *Vref*_*max*_, the reference voltage was manually reduced to *Vref*_*max*_ for safety. Otherwise, the system-determined *Vref*_*opt*_ was used throughout the imaging session.

**Figure 9.**
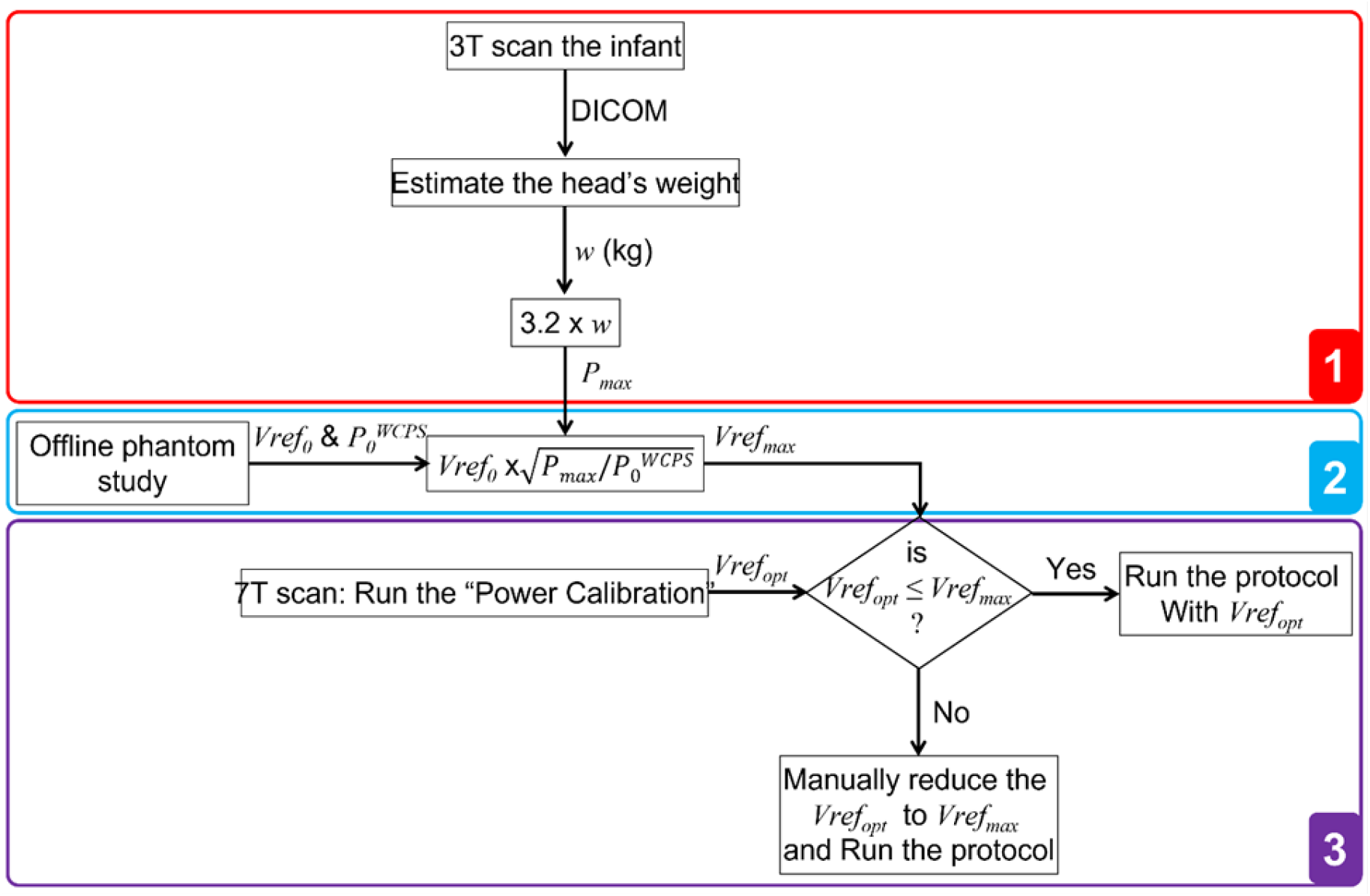
safety workflow combining information from infant 3T scan with offline phantom study to determine the maximum reference voltage for scanning each infant.

#### Infant Imaging

Six infants, weighing between 4.8 kg and 6.1 kg, were scanned at 7T while adhering to the power limits calculated using the workflow described above. Their estimated head weights ranged from 1.08 kg to 1.27 kg, resulting in maximum reference voltages (*Vref*_*max*_) between 181V and 197V for the imaging protocol used in this study (see the Acquisition subsection for protocol details).

To ensure safety and maximal infant comfort during scanning, a temperature probe was placed under the armpit to monitor any changes in core temperature. Consistent with published protocols^48,50,51^ we further equipped infants with multiple layers of hearing protection consisting of moldable silicone ear plugs, held in place by skin sensitive tape and OptoAcoustic headphones (OptoAcoustics Ltd., Tel Aviv). During scanning infants were constantly monitored by a trained staff member inside the magnet room who would stop the scan as soon as infants woke up or showed any other sign of discomfort.

### Acquisition

Data at 3T was acquired at a Siemens Prisma scanner with a 32 channel head coil. fMRI data at 3T was acquired with a four-echo version of the CMRR multiband (MB)-ME sequence^52,53^ (TE = 14 ms, 39 ms, 64 ms, 88 ms, TR = 1.761 s, 2 mm resolution, MB factor = 6, flip angle = 68°). Data at 7T was acquired at a Siemens Magnetom Terra scanner with a one-transmit Nova head coil with 32 receive elements. Functional data at 7T was acquired using two different three-echo sequences (TE = 14 ms, 35 ms, 57 ms, TR = 1.768 s, 1.6 mm resolution, MB factor = 4, flip angle = 60° and TE = 14 ms, 39 ms, 63 ms, TR = 2.39 s, 1.25 mm resolution, MB factor = 3, flip angle = 60°). T_2_w and T_1_w anatomical references were acquired at 3T (T_1_: TR = 2.4 s, TE = 2.2 ms, resolution = 0.8 x 0.8 x 0.8 mm, flip angle = 8°; T_2_: TR = 2.5 s, TE = 323 ms, resolution = 0.8 x 0.8 x 0.8 mm, flip angle = 120° and alternatively T_2_: TR = 4.5 s, TE = 546 ms, resolution = 0.8 x 0.8 x 0.8 mm, flip angle = 120° for PB0021 and PB0022).

During BOLD acquisition (3T and 7T with 1.6 mm sequence) a passive listening auditory paradigm was played to probe event related changes in BOLD activation. A 400 ms long white noise stimulus was played in irregular intervals (9-14 seconds), marking a novel, deviant sound (or oddball) with respect to the regular scanner background noise. Each BOLD scan lasted for 6.7 minutes and contained 24 deviant sounds, starting and ending with ∼1 minute of regular scanner noise.

For PB015 20 such runs were acquired at 3T and 3 at 7T. For PB016 12 runs at 3T and 4 runs at 7T, 13 and 2 for PB017 and 13 and 4 for PB020. For PB017 and PB020, additional short resting state runs with the 7T 1.25 mm sequence were acquired. For PB021 6 runs were acquired at 3T and 4 at 7T and 6 and 11 for PB022 of which 2 runs at 3T and 2 runs at 7T were excluded from analyses due to excessive motion.

### Preprocessing

Data were converted to BIDS format with dcm2bids^54^. Each echo of the ME echo data was denoised with NORDIC^42–44^. Phase and magnitude images of the scan were used for NORDIC as well as three noise frames acquired at the end of each functional run to help estimate the empirical thermal noise level. NORDIC was implemented in Matlab R2019a. Brain tissue segmentations of the anatomical data were generated using BIBSNet^55^, a deep learning model trained on hand edited segmentations of 0-8 month old infants. All data were preprocessed using NiBabies 25.0.1^56^. The options “--multi-step-reg” and “--norm-csf” were used to facilitate registration of native space anatomical data to standard space by including an intermediate age-matched target in template space and M-CRIB-S ^57^ was used for surface reconstruction (“--surface-recon-method mcribs”). For functional data processing the option “--project-goodvoxels” was enabled to exclude voxels with locally high coefficient of variation from volume-to-surface projection of the BOLD time series, and “--cifti-output 91k” to enable output of BOLD time series and morphometric data (surface curvature, sulcal depth, and cortical thickness maps) in the HCP grayordinates space^58^. Susceptibility distortion correction of BOLD time series was done within NiBabies, using SDCFlows with an FSL topup-based method to estimate fieldmaps from “PEPolar” acquisitions (acquisitions with opposite phase encoding direction) ^59^. Data functional connectivity postprocessing was performed using XCP-D 0.10.5^60^ with the “hbcd” mode for setting defaults. Additional options used were “--lower-bpf 0.009” (instead of the default 0.01) to set the lower limit of the band-pass filter (upper limit stayed at the default of 0.08) and “-r 45” to set the head radius. The “--motion-filter-type” - determining range of a band-stop filter - was set to “none” as the TR of the scanning sequences used here were too slow to adequately resolve infants’ fast breathing rate (30-60bpm).

### Data Analysis

T_2_^*^ maps were estimated within the NiBabies preprocessing pipeline using an implementation of Tedana^61^. For quantifying T_2_^*^ relaxation times of cortical vertices, we extracted values from the cortical ribbon and projected them to surface space.

Spatial precision was quantified by estimating FWHM from preprocessed BOLD runs either in native or in MNI space using 3dFWHMx (options -combine -detrend -automask -acf) in AFNI (version 16.1.13)^62^.

Dense connectivity matrices for each subject were constructed from all their low motion (framewise displacement < 0.3mm) 3T or 7T data. A smoothing kernel with sigma 2.25 was used for all matrices.

Stability was quantified as the grayordinate-by-grayordinate correlation of dense functional connectivity matrices within subject similar to Moser et al.^7^ and Lynch et al.^41^. To match stability calculations between 7T and 3T, as available data quantity was very different, this analysis only included 4-5 consecutive runs from each modality (which was the common maximum) compared to all the available data. For stability heat maps, data was split into individual minutes of low motion data and dense connectivity matrices were correlated with each other. Each square on the heatmap represents the average across all included grayordinates. To ensure that results are not biased by natural variations in data quality across multiple runs, the analysis was run with 100 permutations, splitting the whole time series into individual one minute segments which were then shuffled. All results represent the average across these 100 permutations.

## Data and Code Availability

Data can be made available upon request, given a formal data sharing agreement is set up by the institutions involved.

Code is available from the following repository: https://github.com/DCAN-Labs/infant_3T_7T_precision_imaging/

## Author Contributions

JM: Writing - Original Draft, Conceptualization, Investigation, Formal analysis, Visualization, Funding acquisition; AST: Writing - Original Draft, Methodology, Software; JSBR: Formal analysis, Writing - Review & Editing; TJM: Methodology; LH: Project administration, Data Curation; KBW: Methodology, Investigation; HHNP: Data Curation; JTL: Methodology; SS: Investigation; IU: Investigation; ED: Investigation, Project administration; AKL: Writing - Review & Editing; EJA: Writing - Review & Editing; GA: Writing - Review & Editing; CMS: Writing - Review & Editing; SMN: Writing - Review & Editing; YE: Supervision, Resources, Writing - Review & Editing; JTE: Conceptualization, Supervision, Funding acquisition; DAF: Supervision, Funding acquisition; EY: Conceptualization, Investigation, Supervision, Writing - Review & Editing

## Acknowledgements

We thank all the families of our precision babies for their participation and their dedication towards research. We also thank Nick Yamasaki, Lynn Nguyen, Isabella Linder, Angel Gathumbi and Katelyn Day for their support during data acquisition and the Department of Obstetrics, Gynecology and Women’s Health’s Research Administration team for their support with recruitment. The authors acknowledge the Minnesota Supercomputing Institute (MSI) at the University of Minnesota for providing resources that contributed to the research results reported within this paper. URL: http://www.msi.umn.edu.

