## Supplementary Material for "Precision functional imaging in infants using multi-echo fMRI at 7T"

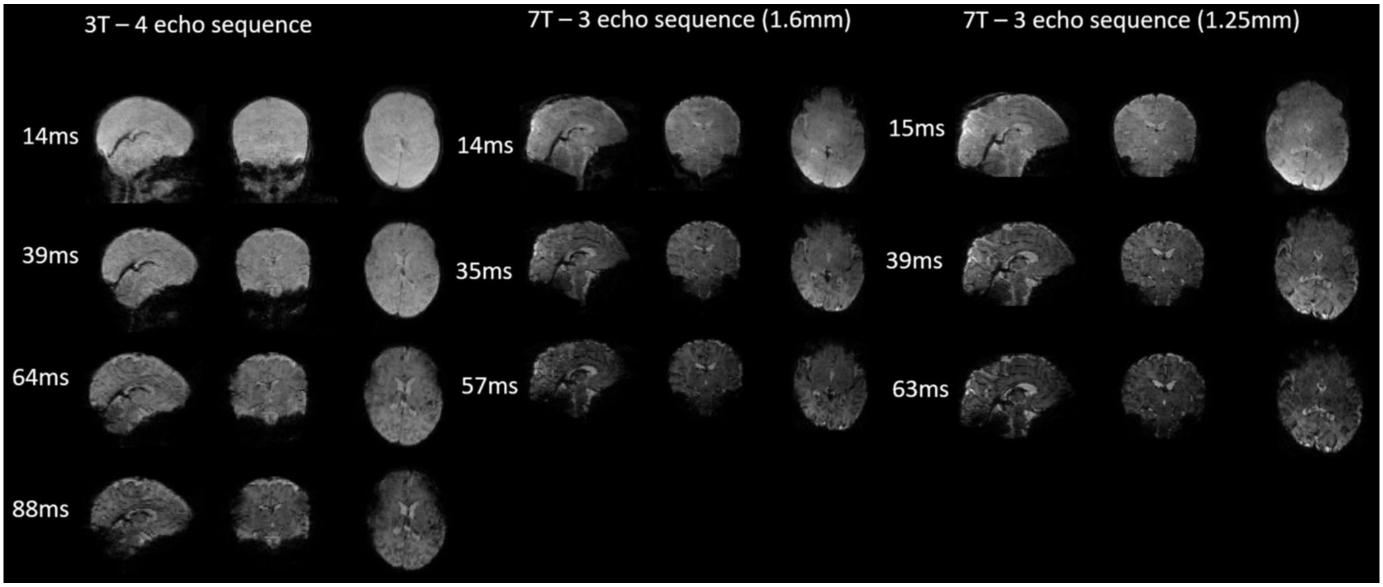


*Suppl. Figure 1: raw multi-echo images from 7T recordings show high spatial tissue contrast. Raw images from the four-echo 3T sequence and the two different three-echo 7T sequences within PB0020.*

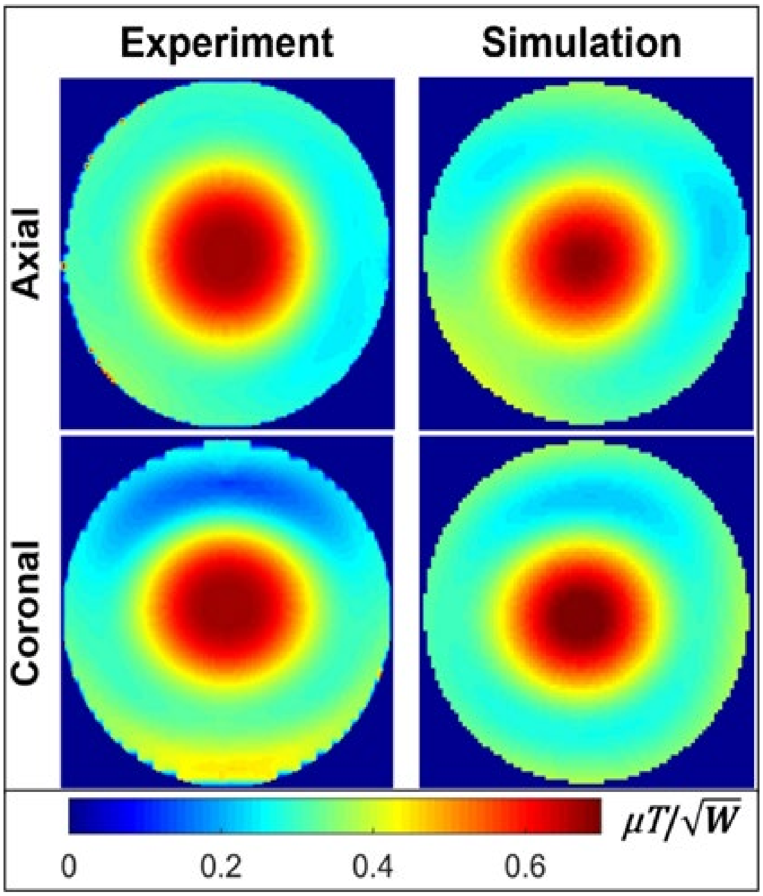


*Suppl. Figure 2: Simulated and measured $B_1^+$ maps*

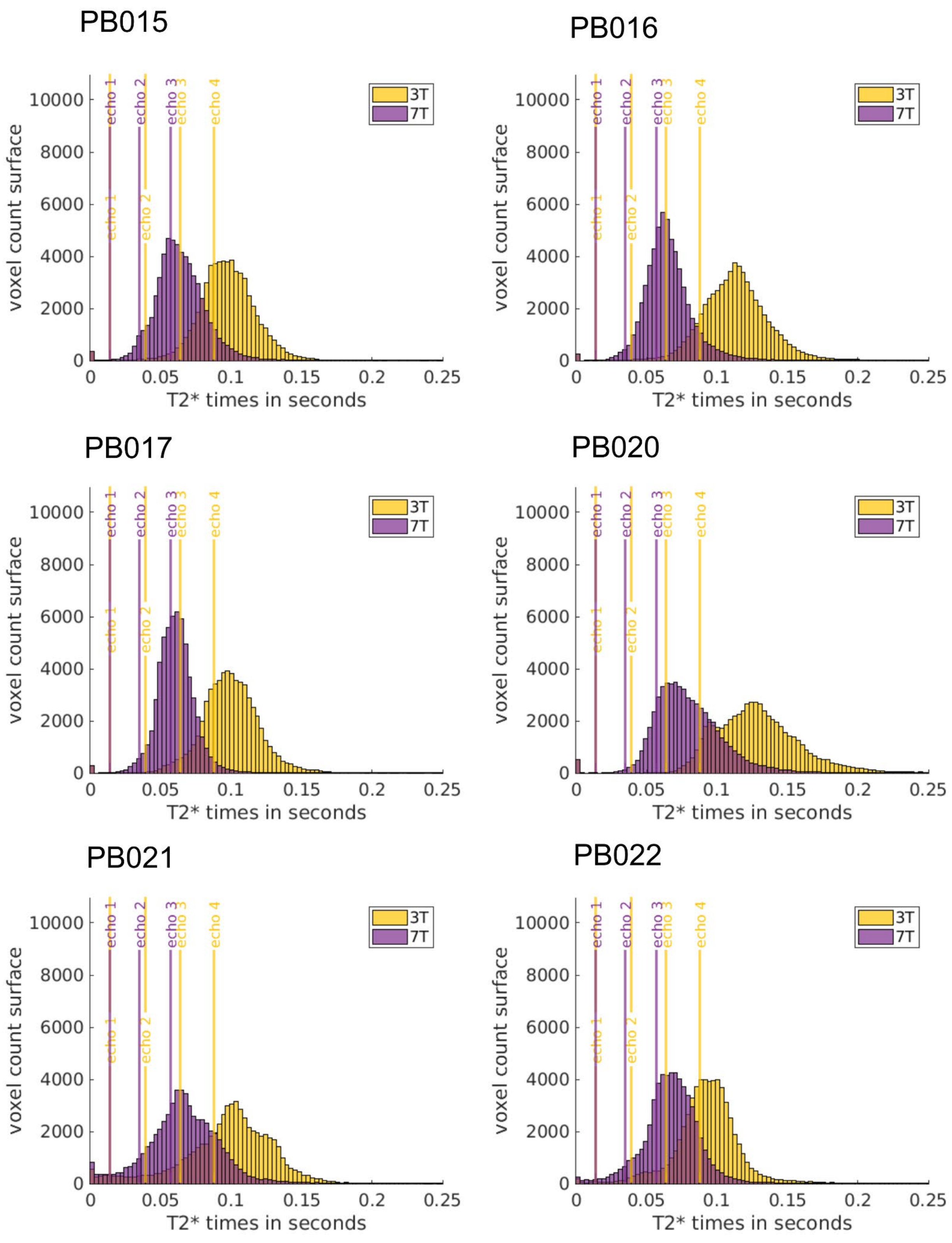


*Suppl Figure 3: distribution of $T_2$* decay times across all cortical grayordinates for each of the participants at 3T and 7T.*

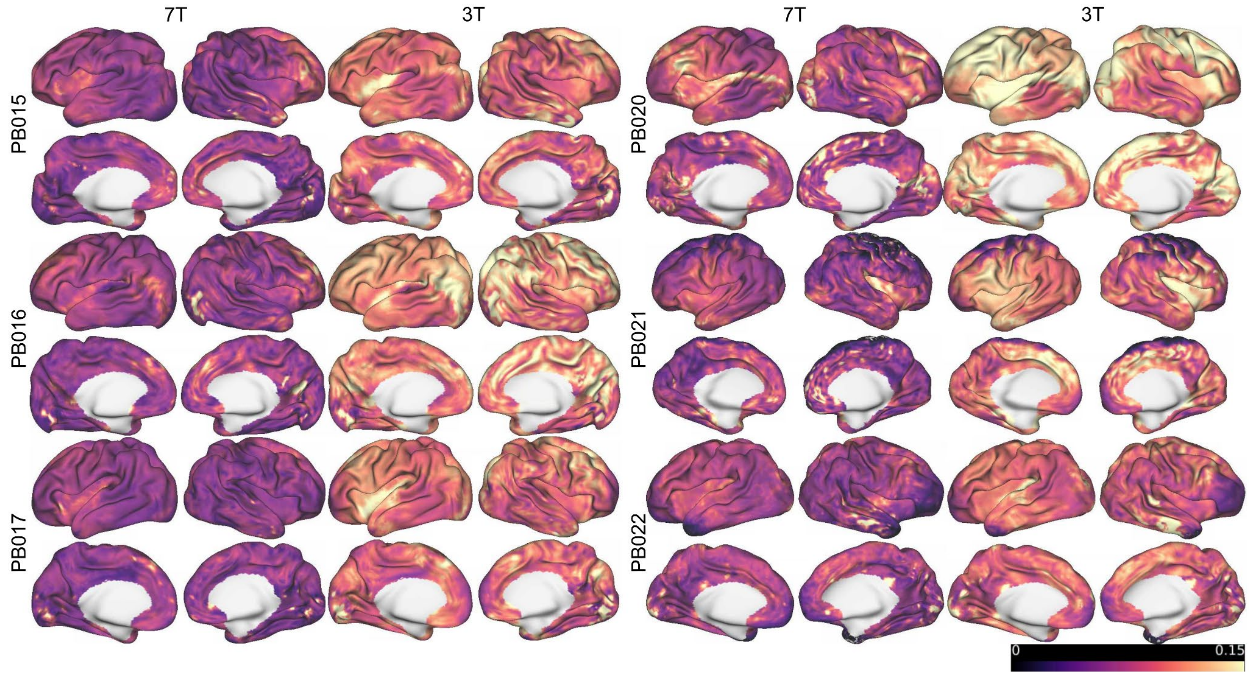


*Suppl Figure 4: Distribution of $T_2$* values across the cortex in 7T and 3T data of the same participants.*

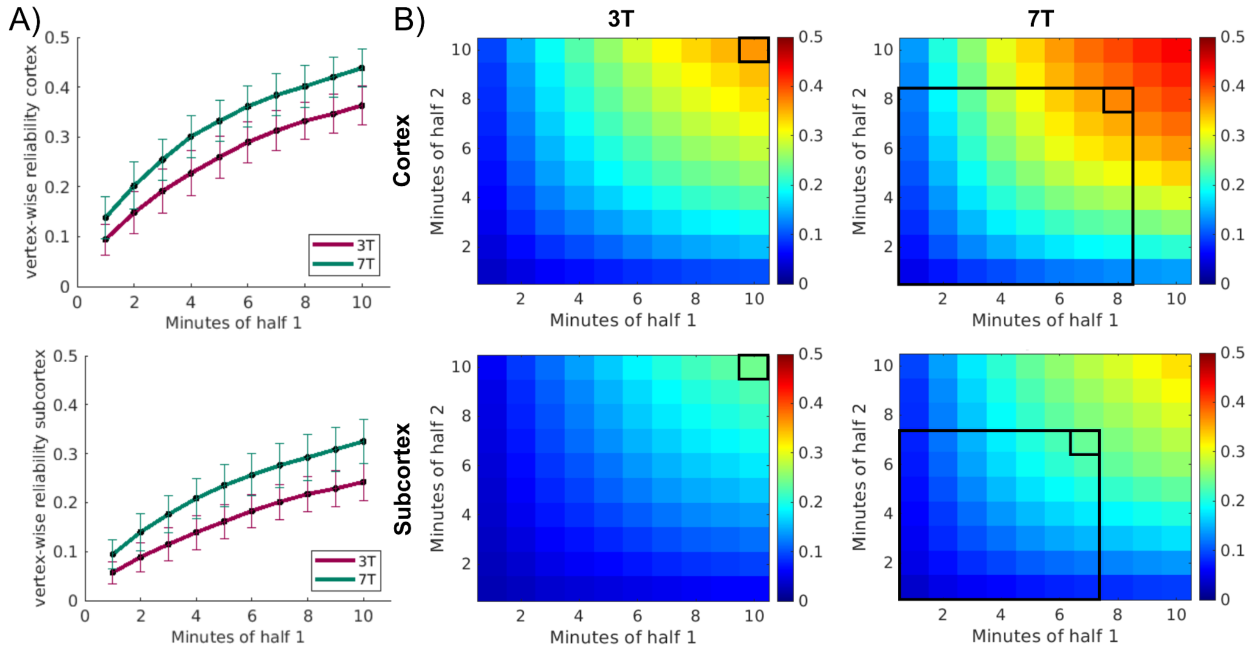


*Suppl Figure 5: Stability heatmap PB020. A) reliability curves for cortical and subcortical vertices. Lines represent the average across vertices and permutations (100 permutations shuffling individual data minutes). Error bars represent the standard deviation across permutations. (Results correspond to the top row of heatmaps in B)) B) Each square represents the correlation of connectivity matrices when a set amount of data is used to create them (extending Figure A to also varying data amounts in half 2). Correlation strength is represented by the heatmap. Values are averaged across vertices and permutations. Rectangles indicate the amount of 7T data that is equivalent to the maximum in the 3T data.*

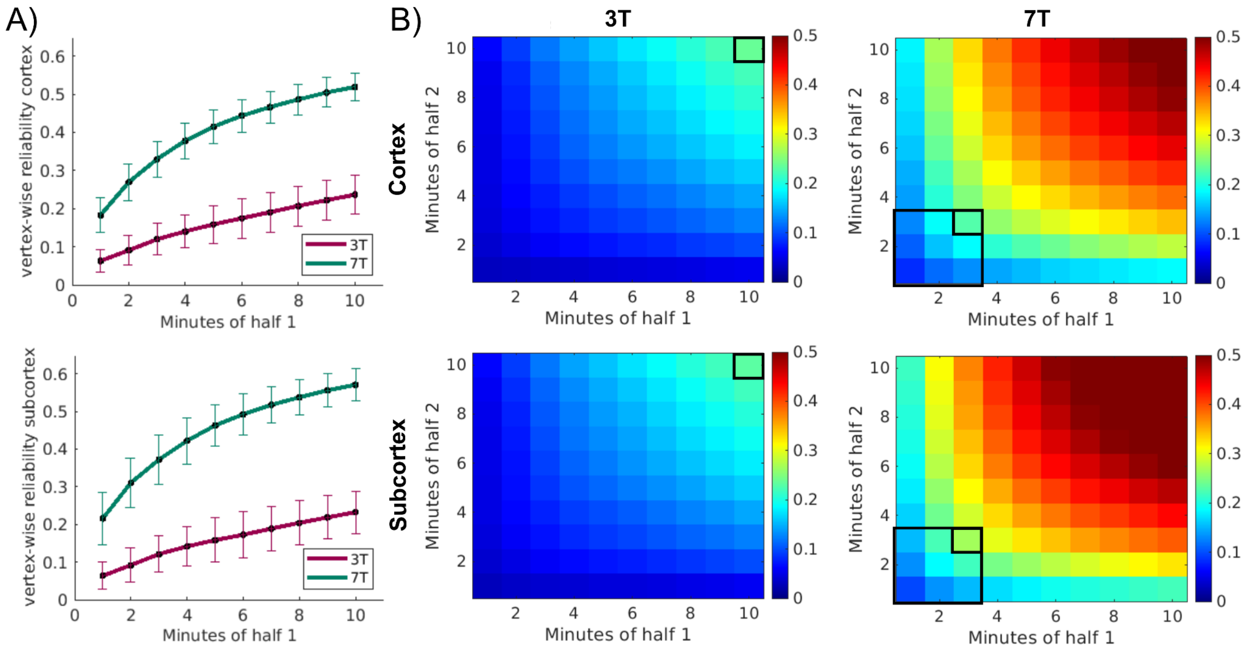


*Suppl Figure 6: Stability heatmap PB021. A) reliability curves for cortical and subcortical vertices. Lines represent the average across vertices and permutations (100 permutations shuffling individual data minutes). Error bars represent the standard deviation across permutations. (Results correspond to the top row of heatmaps in B)) B) Each square represents the correlation of connectivity matrices when a set amount of data is used to create them (extending Figure A to also varying data amounts in half 2). Correlation strength is represented by the heatmap. Values are averaged across vertices and permutations. Rectangles indicate the amount of 7T data that is equivalent to the maximum in the 3T data.*

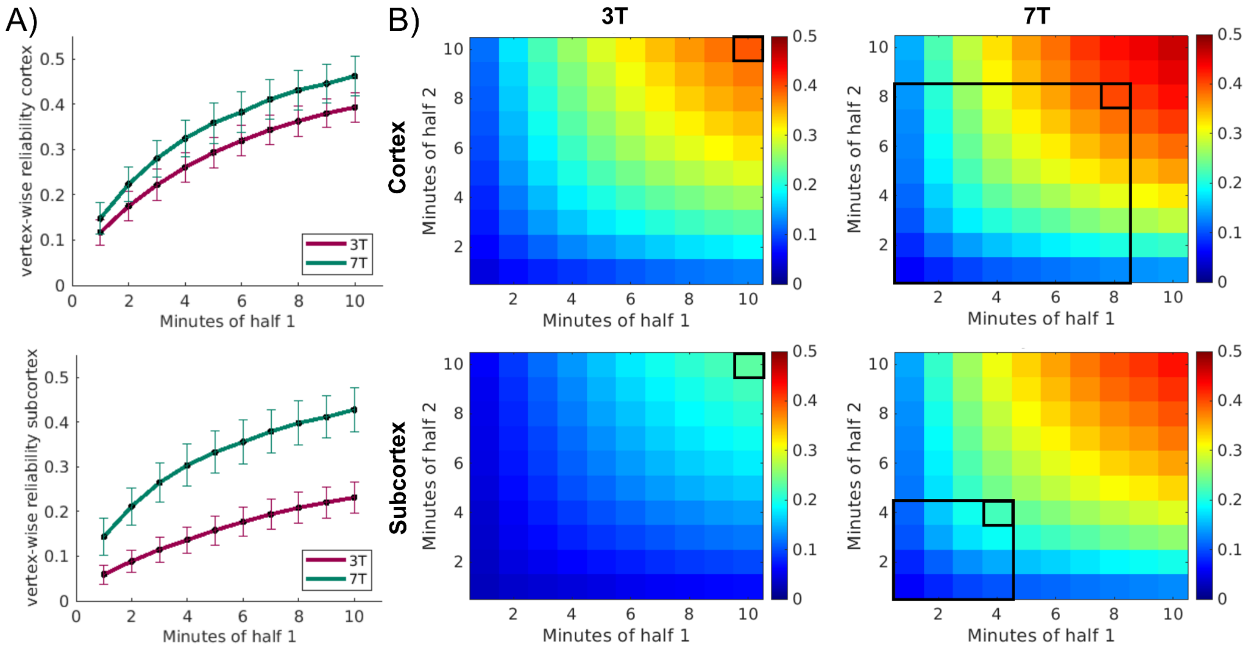


*Suppl Figure 7: Stability heatmap PB022. A) reliability curves for cortical and subcortical vertices. Lines represent the average across vertices and permutations (100 permutations shuffling individual data minutes). Error bars represent the standard deviation across permutations. (Results correspond to the top row of heatmaps in B)) B) Each square represents the correlation of connectivity matrices when a set amount of data is used to create them (extending Figure A to also varying data amounts in half 2). Correlation strength is represented by the heatmap. Values are averaged across vertices and permutations. Rectangles indicate the amount of 7T data that is equivalent to the maximum in the 3T data.*

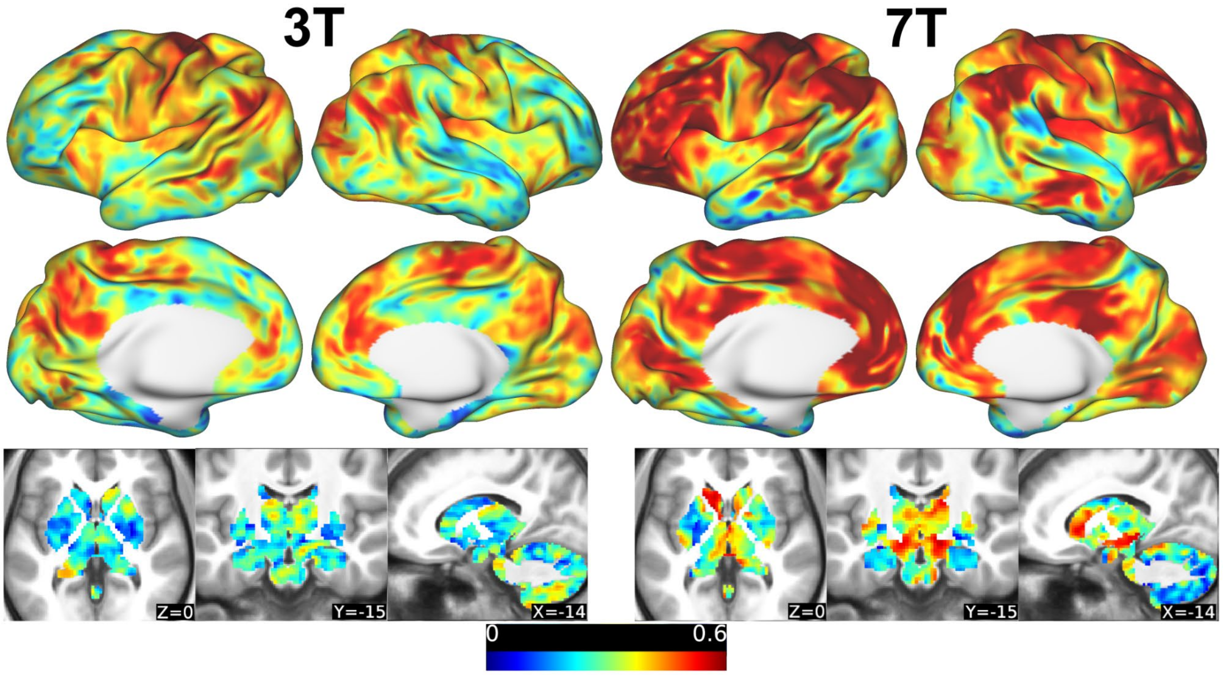


*Suppl Figure 8: Spatial distribution of stability map from PB020 using 10 min vs. 10 min of low motion data (top right corner Suppl Figure 5 A and B) for 3T and 7T. Maps show common regions of low stability and regions of particularly high stability in 7T data.*

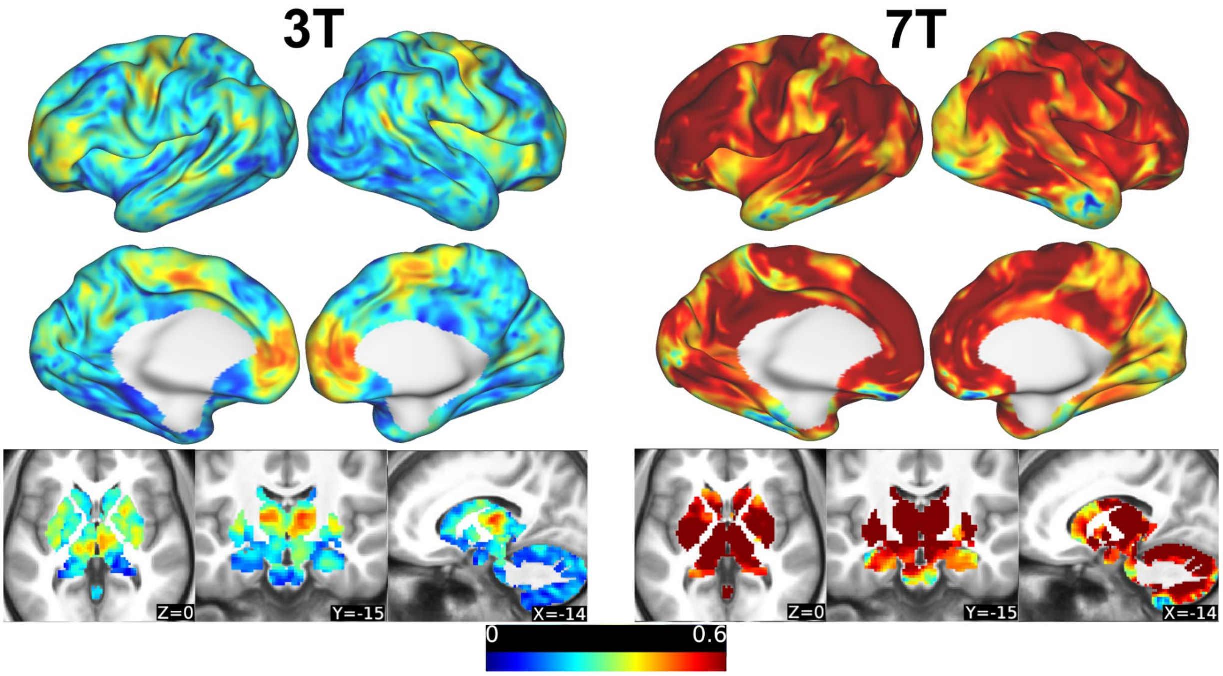


*Suppl Figure 9: Spatial distribution of stability map from PB021 using 10 min vs. 10 min of low motion data (top right corner Suppl Figure 6 A and B) for 3T and 7T. Maps show common regions of low stability and regions of particularly high stability in 7T data.*

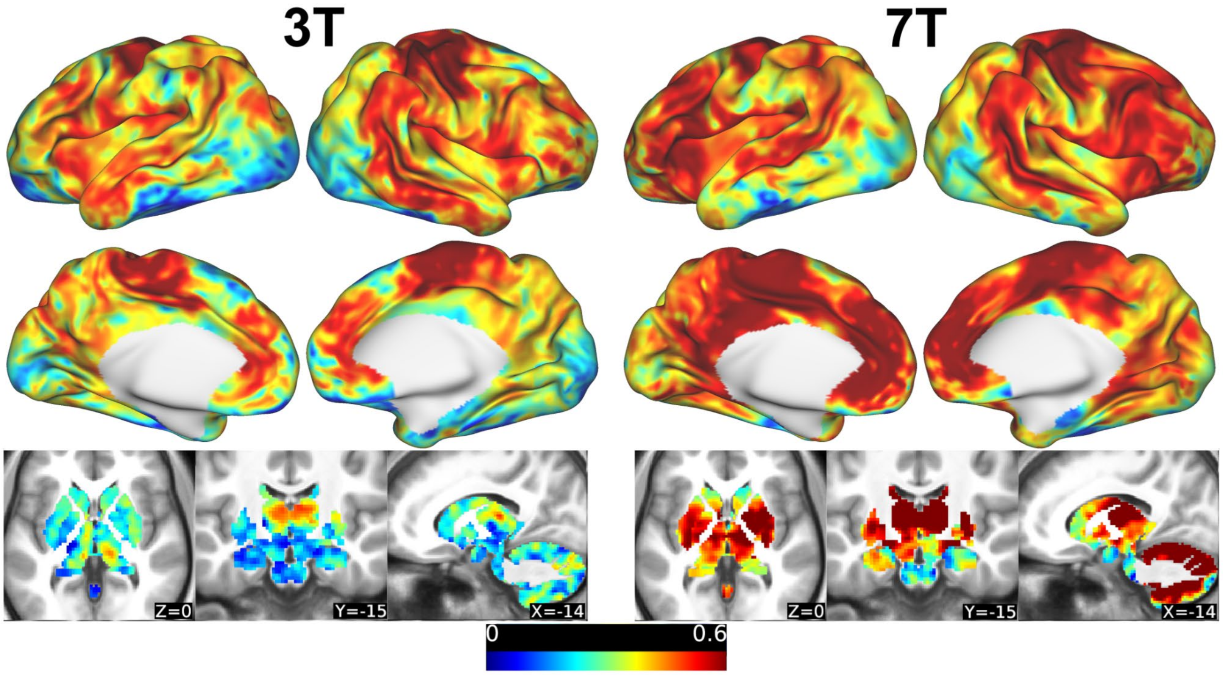


*Suppl Figure 10: Spatial distribution of stability map from PB022 using 10 min vs. 10 min of low motion data (top right corner Suppl Figure 7 A and B) for 3T and 7T. Maps show common regions of low stability and regions of particularly high stability in 7T data.*

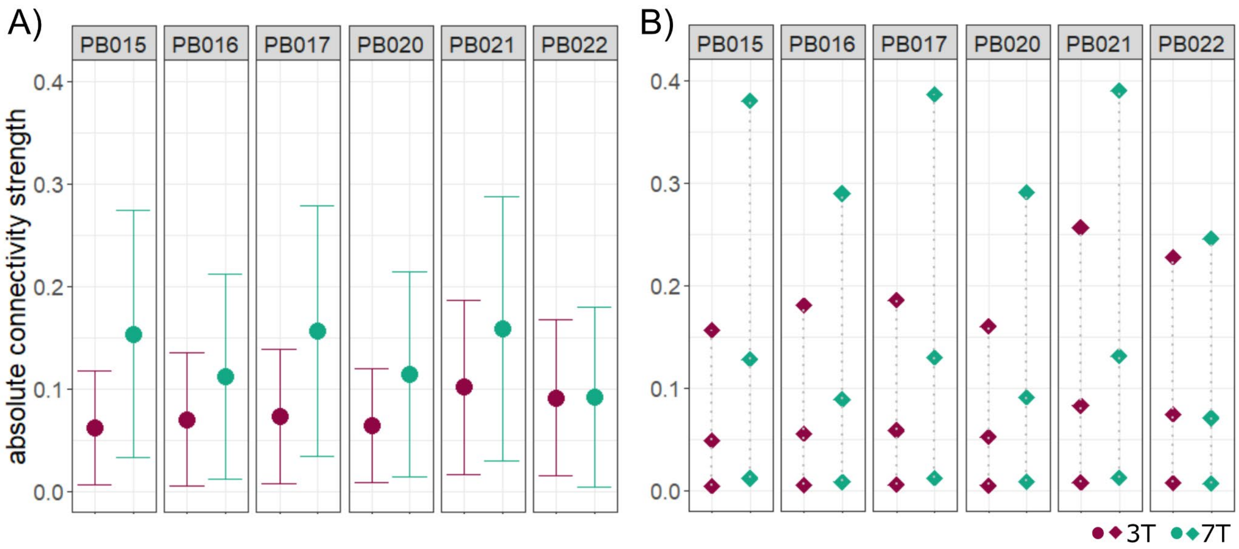


*Suppl Figure 11: Absolute strength of functional connections of subcortical grayordinates compared between 3T and 7T data of each participant. A) mean with standard deviation of absolute connections B) 5th, 50th and 95th percentiles of absolute connections. All metrics show stronger or equivalent connectivity for 7T.*

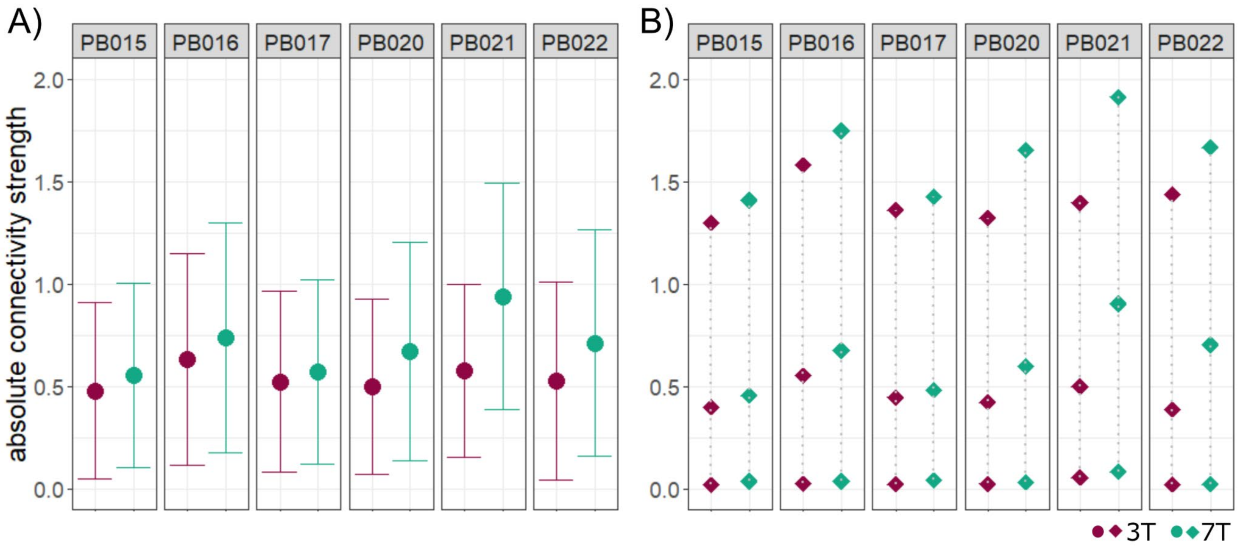


*Suppl Figure 12: Absolute strength of functional connections of short range connections (radius <1 cm) compared between 3T and 7T data of each participant. A) mean with standard deviation of absolute connections B) 5th, 50th and 95th percentiles of absolute connections. All metrics show stronger or equivalent connectivity for 7T.*

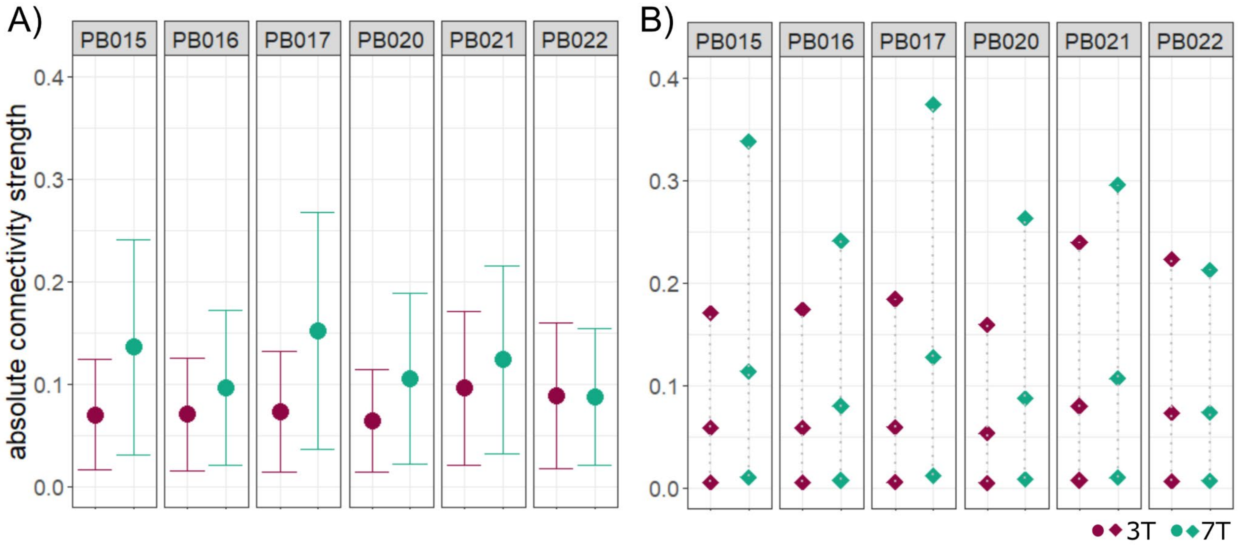


*Suppl Figure 13: Absolute strength of functional connections of long range connections (further than 10 cm from seed) compared between 3T and 7T data of each participant. A) mean with standard deviation of absolute connections B) 5th, 50th and 95th percentiles of absolute connections. All metrics show stronger or equivalent connectivity for 7T.*

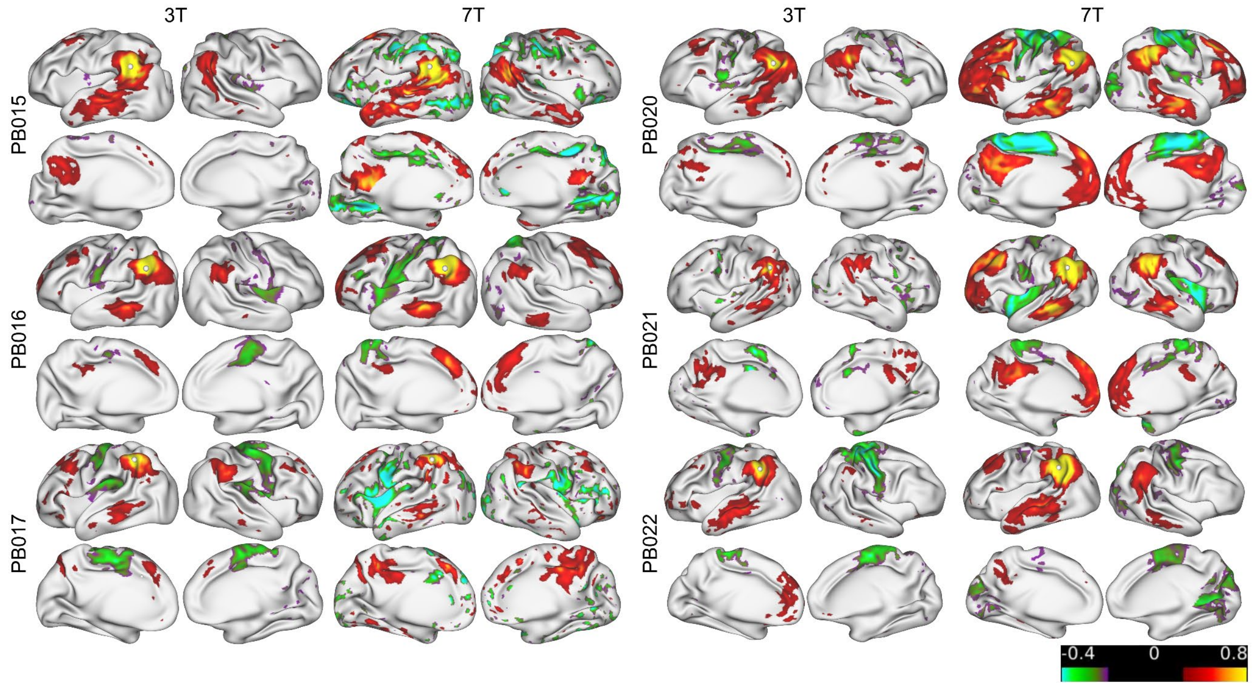


*Suppl Figure 14: Example DMN connectivity for all participants. In seed maps from 7T data within network correlations between anterior and posterior hubs and the contralateral hemisphere are stronger as well as characteristic negative correlations with for example the motor cortex. The seed used here is marked by a white dot.*

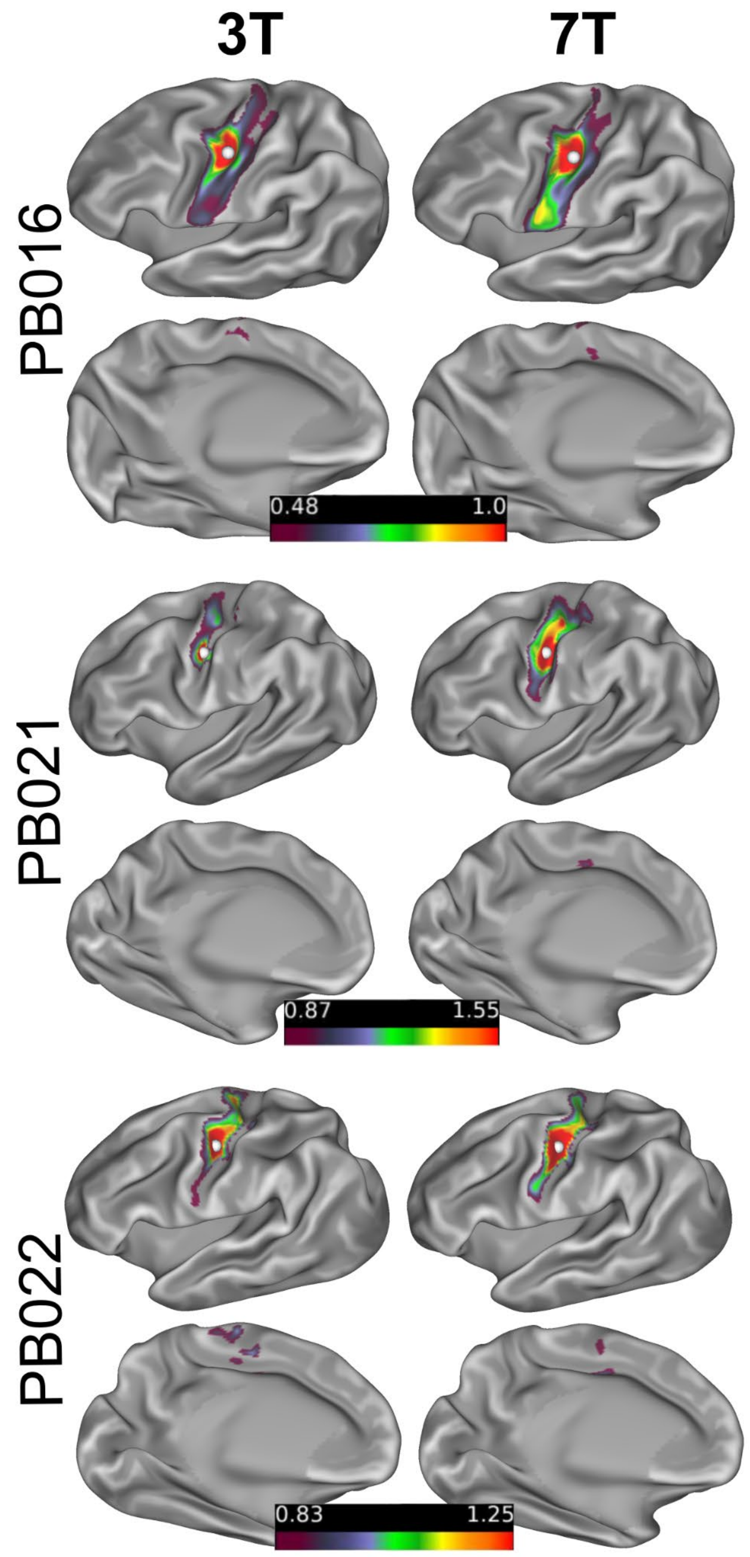


*Suppl Figure 15: Example SCAN in 3T and 7T data for the three other participants with over 20 minutes of 7T data (all 7T data are 1.6 mm). Seed is placed in the middle SCAN hub.*
